# Ebselen derivatives are very potent dual inhibitors of SARS-CoV-2 proteases - PL^pro^ and M^pro^ in in vitro studies

**DOI:** 10.1101/2020.08.30.273979

**Authors:** Mikolaj Zmudzinski, Wioletta Rut, Kamila Olech, Jarosław Granda, Mirosław Giurg, Małgorzata Burda-Grabowska, Linlin Zhang, Xinyuanyuan Sun, Zongyang Lv, Digant Nayak, Malgorzata Kesik-Brodacka, Shaun K. Olsen, Rolf Hilgenfeld, Marcin Drag

**Affiliations:** Department of Chemical Biology and Bioimaging, Wroclaw University of Science and Technology, Wyb. Wyspianskiego 27, 50-370 Wroclaw, Poland; Department of Organic and Medicinal Chemistry, Faculty of Chemistry, Wroclaw University of Science and Technology, Wyb. Wyspianskiego 27, 50-370 Wrocław, Poland; Institute of Molecular Medicine, University of Lübeck, Ratzeburger Allee 160, 23562 Lübeck, Germany; Institute of Biochemistry, Center for Structural and Cell Biology in Medicine, University of Lübeck, Ratzeburger Allee 160, 23562 Lübeck, Germany; German Center for Infection Research (DZIF), Hamburg-Lübeck-Borstel-Riems Site, University of Lübeck, 23562 Lübeck, Germany; Department of Biochemistry & Structural Biology University of Texas Health Science Center at San Antonio, San Antonio, TX, 78229 USA; Research Network Łukasiewicz - Institute of Biotechnology and Antibiotics, Starościńska 5, 02-516, Warsaw, Poland; Center for Brain, Behavior and Metabolomics (CBBM), University of Lübeck, 23562 Lübeck, Germany

**Keywords:** Coronavirus, organoselenium compounds, COVID-19, cysteine protease

## Abstract

Proteases encoded by SARS-CoV-2 constitute a promising target for new therapies against COVID-19. SARS-CoV-2 main protease (M^pro^, 3CL^pro^) and papain-like protease (PL^pro^) are responsible for viral polyprotein cleavage - a process crucial for viral survival and replication. Recently it was shown that 2-phenylbenzisoselenazol-3(2H)-one (ebselen), an organoselenium anti-inflammatory small-molecule drug, is a potent, covalent inhibitor of both the proteases and its potency was evaluated in enzymatic and anti-viral assays. In this study, we screened a collection of 23 ebselen derivatives for SARS-CoV-2 PL^pro^ and M^pro^ inhibitors. Our studies revealed that ebselen derivatives are potent inhibitors of both the proteases. We identified three PL^pro^ and four M^pro^ inhibitors superior to ebselen. Our work shows that ebselen constitutes a promising platform for development of new antiviral agents targeting both SARS-CoV-2 PL^pro^ and M^pro^.

## 1. Introduction

In the winter of 2019, an outbreak of pneumonia with flu-like symptoms emerged in Wuhan, China.^1,2^ Shortly thereafter, the disease-causing pathogen was isolated and analyzed, leading to identification of the novel, highly contagious human beta-coronavirus SARS-CoV-2 (formerly known as 2019-nCoV).^3^ By the end of August 2020, with over 24.5 million people diagnosed with Coronavirus Disease 2019 (COVID-19), the death toll exceeded 830,000 patients worldwide.^4^ With neither vaccines nor drugs targeting the virus available, various strategies have been employed to accelerate finding an effective therapy to fight the pathogen.^5^ One of these strategies is drug repurposing - establishing therapeutic properties for already approved substances for new medical applications. This strategy can be supported by computational analysis, which can lower the costs, speed up the process in comparison with *de-novo* development of new therapeutics and serve as a first stage in screening vast libraries of active compound.^6–10^ Drug repositioning has been already successfully used in fighting COVID-19.^11^ A bright example here is remdesivir, an antiviral agent targeting viral RNA-dependent RNA polymerase (RdRp) that was designated to treat Ebola but has shown efficacy shortening recovery time and reducing mortality as well as serious adverse effects in COVID-19 patients.^12^ Nonetheless, current treatment options are critically limited and finding new therapeutics for COVID-19 patients constitutes a leading challenge for the scientific community.

To address the problem, medicinal chemists identified druggable targets among viral non-structural proteins (nsps), two of them being proteases. The SARS-CoV-2 main protease (M^pro^, 3CL^pro^, nsp5) and the papain-like protease (PL^pro^, nsp3 papain-like protease domain) enable viral replication in host cells by processing the viral polyprotein and generating 16 nsps, crucial for virus replication. SARS-CoV-2 M^pro^ generates 13 viral nsps, making it a key player in the process of virus replication and maturation.^13–15^ M^pro^ is a dimeric cysteine protease with a structure highly conserved among human coronaviruses. Unusual preference for a glutamine residue at the P1 position of the substrate cleavage site sets M^pro^ apart from known human proteases. This feature can be beneficial for design and synthesis of effective, broad-spectrum antiviral agents with minimum side effects.^7,13,16–19^ SARS-CoV-2 PL^pro^ is a viral cysteine protease proposed as an excellent target for COVID-19 treatment due to its pathophysiological roles. PL^pro^ processes viral polyprotein and generates proteins nsp1-3. Moreover, the protease also alters the host immune response by deubiquitinating and deISGylating proteins within infected cells.^20–23^ Thus, PL^pro^ inhibition would not only block the replication of the virus, but would also limit the dysregulation of cellular signaling mediated by ISG15 and ubiquitin.

2-phenylbenzisoselenazol-3(2H)-one (ebselen) is a small-molecule drug with a pleiotropic mode of action in cells.^24^ Ebselen is an excellent scavenger of ROS that acts as a mimic of glutathione peroxidase (GPx) and interacts with the thioredoxin (Trx) system by oxidation of reduced TrxR.^25–27^ Recently it was shown that ebselen inhibits both the SARS-CoV-2 proteases. Weglarz-Tomczyk et al. evaluated ebselen^28^ and a collection of its derivatives^29^ as inhibitors of PL^pro^, leading to identification of inhibitors with IC_50_ values in the nanomolar range. In another study, a library of approx. 10,000 drugs and drug candidates was screened for M^pro^ inhibitors. As a result, ebselen displayed the lowest IC_50_ among the substances tested (0.67 μM), furthermore it also displayed an antiviral effect in SARS-CoV-2 infected Vero cells.^7^

In this work, we used ebselen and a collection of 23 of its derivatives to evaluate their properties as SARS-CoV-2 PL^pro^ and M^pro^ inhibitors. First, we screened the collection for inhibitors of both proteases. Next, we determined the half-maximum inhibitory concentration (IC_50_) values for the most promising hits. We show that ebselen may constitute a potential lead compound for development of novel antiviral agents. The results can be useful in the design of new active compounds targeting the proteases encoded by SARS-CoV-2, to be applied in COVID-19 treatment.

## 2. Results & Discussion

The efficacy of ebselen and other organoselenium compounds has been previously evaluated for HIV^30,31^, HSV2^32^, HCV^33^, and Zika virus^34^ infections. Moreover, a recent report presents ebselen and its derivatives as potent inhibitors of SARS-CoV-2 PL^pro.29^ In order to find new inhibitors of proteases encoded by the new coronavirus, we screened a collection of ebselen and its 23 derivatives with mono- or disubstitutions within the phenyl ring (Tab. 1).

### 2.1. Compound library screening for SARS-CoV-2 PL^pro^ inhibitors

First, we evaluated the inhibitory properties for the compounds at 1 μM inhibitor concentration and 100 nM SARS-CoV-2 PL^pro^. For the assay, we used the fluorogenic substrate Ac-LRGG-ACC with a structure based on the C-terminal epitope of Ub and ISG15 proteins as well as on the nsp1/2, nsp2/3, and nsp3/4 cleavage sites in the coronaviral polyprotein. PL^pro^ screening resulted in identification of only one compound (**7**) with higher potency (84.4%) than ebselen (65.4%). However, **7** differed from the other compounds in the collection as it was the only investigated ebselen derivative with a 3-substituted pyridinyl moiety instead of a substituted phenyl ring. The results also show that electron-withdrawing groups (EWGs) at the ortho position of the phenyl ring hamper inhibition of the PL^pro^ by the compounds. Derivatives with strong EWGs (trifluromethyl group for **5** and nitro group for **6**) displayed the 2^nd^ and 3^rd^ lowest potency, while the other compounds displayed approx. 50% of PL^pro^ inhibition. We did not observe any significant differences between inhibitory properties between mono- and disubstituted ebselen derivatives.

### 2.2. Compound library screening for M^pro^ inhibitors

Next, we screened the library at 100 nM inhibitors and 100 nM M^pro^ concentrations. For the assay, we used a novel tetrapeptide fluorogenic substrate for SARS-CoV-2 M^pro^, QS1 (Ac-Abu-Tle-Leu-Gln-ACC; K_M_=207.3±12 μM, k_cat_/K_M_=859±57 M^-1^s^-1^).^17^ As a result, ebselen displayed 57.6% M^pro^ inhibition. The best hits were compounds **10** and **17** with >80% of M^pro^ inhibition. **10** represents monosubstituted derivatives with a nitro group at the *para* position. Other substitutions at the *para* position do not seem to have such a strong effect on inhibition of M^pro^. On the other hand, our second best hit, **17**, has a 5-chloro-2-fluoro disubstituted phenyl ring and represents a group of ebselen disubstituted derivatives. Analogically to **10**, we did not identify any other 2,5-disubstitutions providing a similar effect on M^pro^ inhibition. Interestingly, the third best inhibitor (77.1%) has a 2,4-disubstituted phenyl ring. We also observed, that 2,4-dimethoxy derivative (**16)** displays potency towards both of the proteases close to ebselen’s, however, in comparison with ebselen, its toxicity evaluated in A549 human cell line was 10 times lower.^35^ In general, substitutions within the phenyl ring of ebselen boost inhibition of M^pro^ as we identified only 3 compounds (**6, 7, 12**) with a potency lower than for ebselen.

**Table 1.**
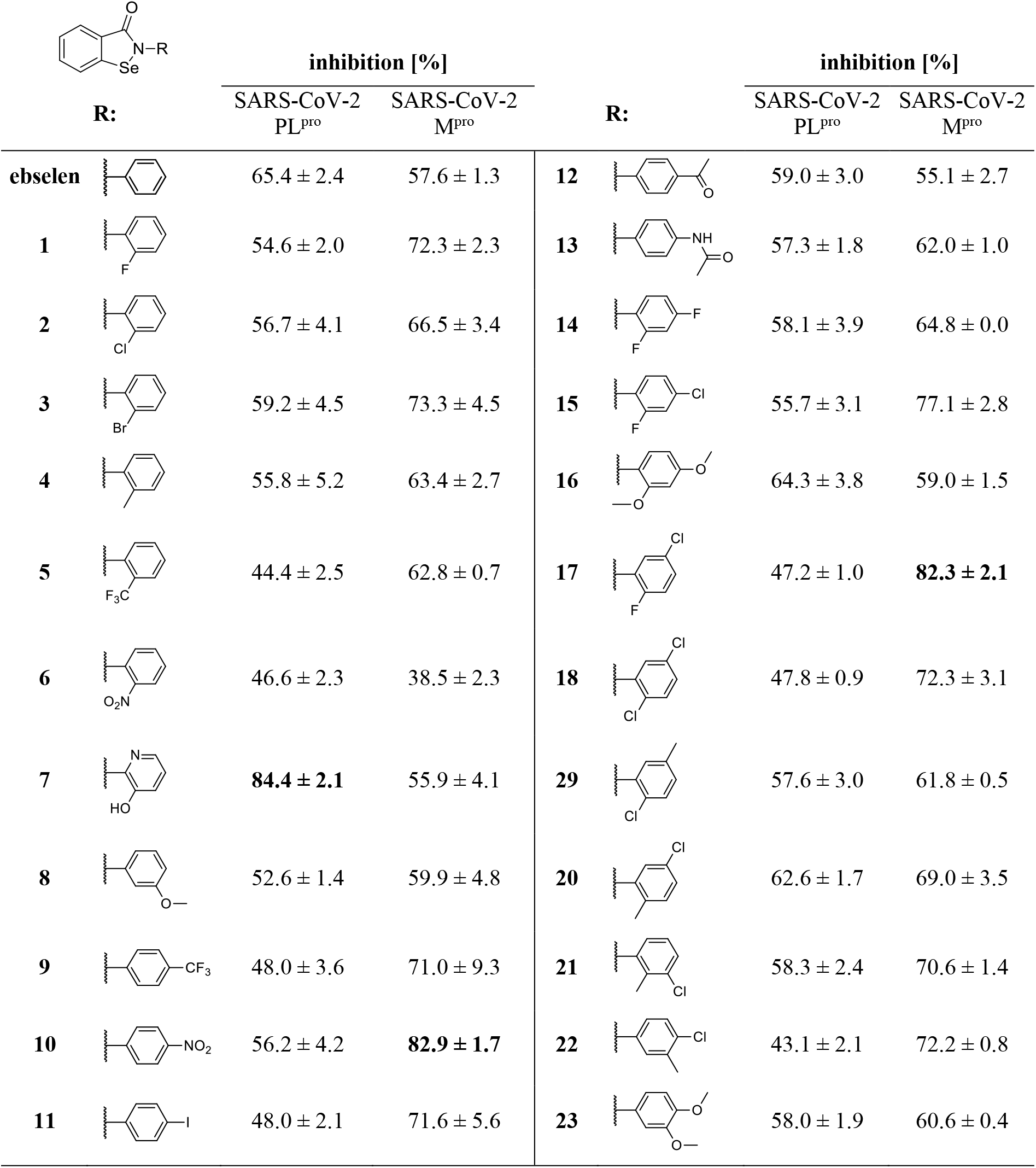
The results of inhibitor screening for SARS-CoV-2 proteases. Assay conditions for SARS-CoV-2 PL^pro^: [E]=100 nM, [I]=1 μM, [S]=10 μM; for SARS-CoV-2 M^pro^: [E]=100 nM, [I]=100 nM, [S]=50 μM.

| **R:** | inhibition [%] |  | **R:** | inhibition [%] |  |
| --- | --- | --- | --- | --- | --- |
|  | SARS-CoV-2<br>PL<sup>pro</sup> | SARS-CoV-2<br>M<sup>pro</sup> |  | SARS-CoV-2<br>PL<sup>pro</sup> | SARS-CoV-2<br>M<sup>pro</sup> |
| **ebselen** | 65.4 ± 2.4 | 57.6 ± 1.3 | **12** | 59.0 ± 3.0 | 55.1 ± 2.7 |
| **1** | 54.6 ± 2.0 | 72.3 ± 2.3 | **13** | 57.3 ± 1.8 | 62.0 ± 1.0 |
| **2** | 56.7 ± 4.1 | 66.5 ± 3.4 | **14** | 58.1 ± 3.9 | 64.8 ± 0.0 |
| **3** | 59.2 ± 4.5 | 73.3 ± 4.5 | **15** | 55.7 ± 3.1 | 77.1 ± 2.8 |

|  |  |  |  |  |  |  |  |
| --- | --- | --- | --- | --- | --- | --- | --- |
| **4** | | $55.8 \pm 5.2$ | $63.4 \pm 2.7$ | **16** | | $64.3 \pm 3.8$ | $59.0 \pm 1.5$ |
| **5** | | $44.4 \pm 2.5$ | $62.8 \pm 0.7$ | **17** | | $47.2 \pm 1.0$ | **$82.3 \pm 2.1$** |
| **6** | | $46.6 \pm 2.3$ | $38.5 \pm 2.3$ | **18** | | $47.8 \pm 0.9$ | $72.3 \pm 3.1$ |
| **7** | | **$84.4 \pm 2.1$** | $55.9 \pm 4.1$ | **29** | | $57.6 \pm 3.0$ | $61.8 \pm 0.5$ |
| **8** | | $52.6 \pm 1.4$ | $59.9 \pm 4.8$ | **20** | | $62.6 \pm 1.7$ | $69.0 \pm 3.5$ |
| **9** | | $48.0 \pm 3.6$ | $71.0 \pm 9.3$ | **21** | | $58.3 \pm 2.4$ | $70.6 \pm 1.4$ |
| **10** | | $56.2 \pm 4.2$ | **$82.9 \pm 1.7$** | **22** | | $43.1 \pm 2.1$ | $72.2 \pm 0.8$ |
| **11** | | $48.0 \pm 2.1$ | $71.6 \pm 5.6$ | **23** | | $58.0 \pm 1.9$ | $60.6 \pm 0.4$ |

### 2.3. IC_50_ determination

Based on the screening results, we selected ebselen and seven of its derivatives for further inhibitory property evaluation. We chose compounds: a), exhibiting the highest potency towards M^pro^ (**10, 17**) or PL^pro^ (**7**) in the screening assay; or b), displaying relatively high inhibition towards both the investigated proteases (**3, 16, 20, 21**) (Fig. 2). During the assays, IC_50_ values for PL^pro^ were in the micromolar range while for M^pro^, they were in the low nanomolar range. For the reference inhibitor, ebselen, IC_50_ values were respectively 1.12μM ± 0.06 and 30.91 ± 2.67 nM. Compound **7**, which was the best hit in the PL^pro^ inhibitor screening assay, indeed had the lowest IC_50_ value (0.58 **±** 0.04 μM) among the tested compounds. Despite ebselen being the second best PL^pro^ inhibitor in the screening assay, we found that two other compounds (**17, 21**) displayed slightly lower IC_50_ values. As expected, **10** and **17**, which were selected for the analysis as the best M^pro^ inhibitors, displayed lower IC_50_ values than ebselen. The most potent M^pro^ inhibitor with IC_50_=15.24 nM was compound **17**, the second best hit from the screening experiment. Interestingly, the best hit (**11**) displayed an IC_50_ value similar to values determined for **3** and **21** (respectively 27.95, 25.69, and 27.37 nM). For **16** and **20**, we observed that despite a higher potency in the screening assay, the IC_50_ values determined for these compounds were higher than for ebselen. In general, the screening results correlated well with the determined IC_50_ values (Tab. 2).

**Figure 2.**
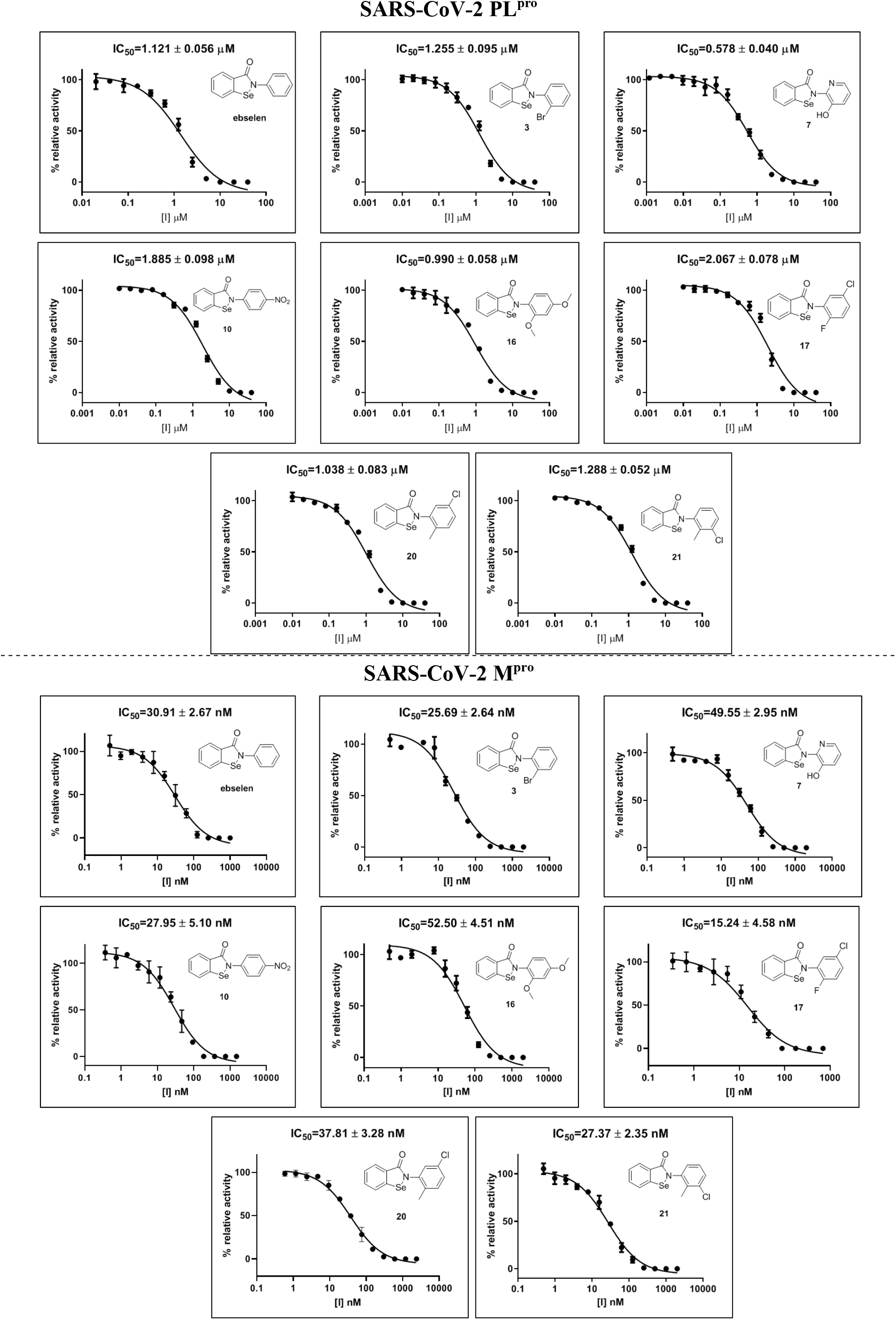
IC_50_ values of SARS-CoV-2 M^pro^ and PL^pro^ inhibitors.

**Table 2.**
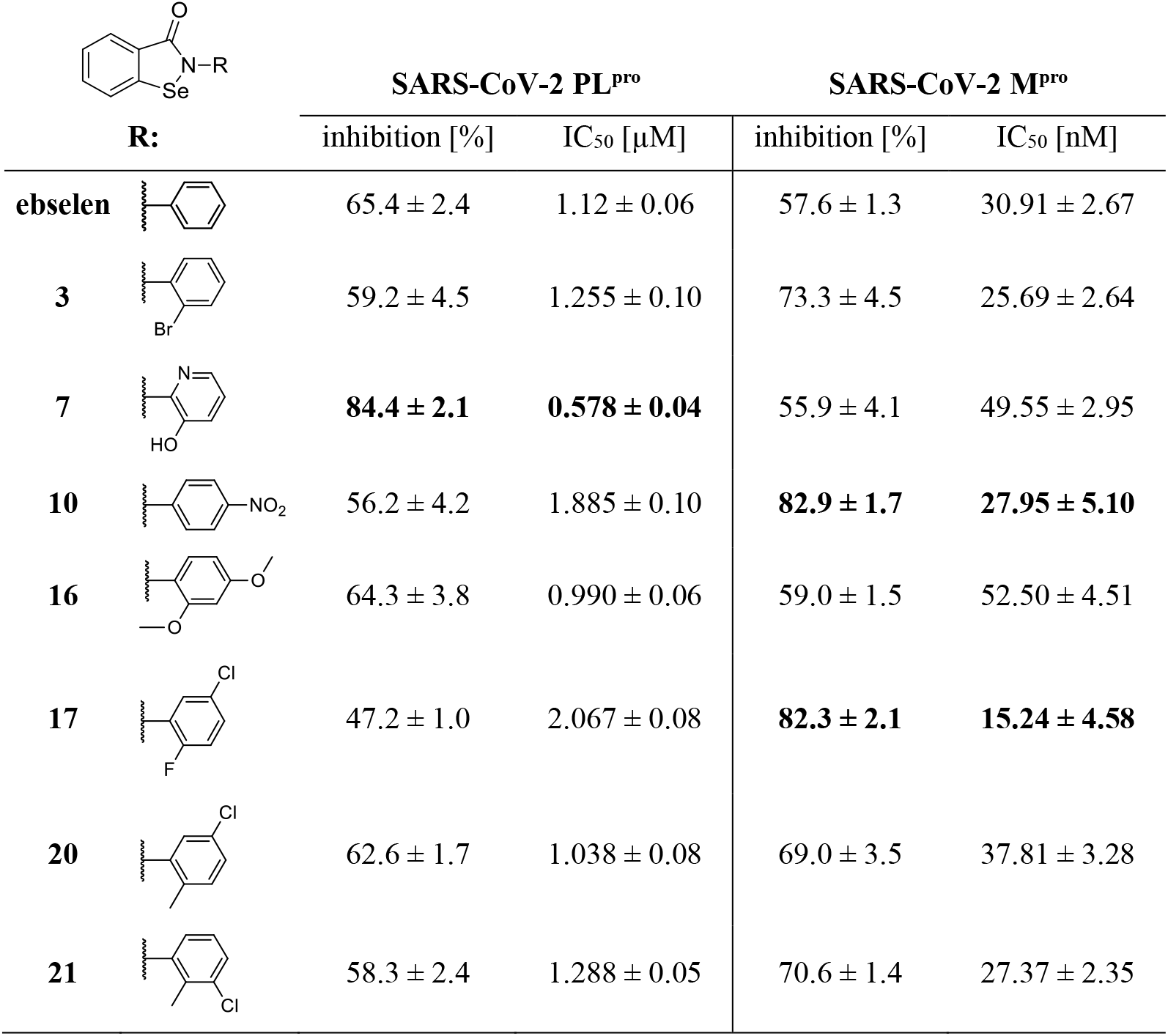
Inhibitory properties for the selected compounds.

| **R:** | **SARS-CoV-2 PL<sup>pro</sup>** |  | **SARS-CoV-2 M<sup>pro</sup>** |  |
| --- | --- | --- | --- | --- |
|  | inhibition [%] | IC<sub>50</sub> [μM] | inhibition [%] | IC<sub>50</sub> [nM] |
| **ebselen** | 65.4 ± 2.4 | 1.12 ± 0.06 | 57.6 ± 1.3 | 30.91 ± 2.67 |
| **3** | 59.2 ± 4.5 | 1.255 ± 0.10 | 73.3 ± 4.5 | 25.69 ± 2.64 |
| **7** | **84.4 ± 2.1** | **0.578 ± 0.04** | 55.9 ± 4.1 | 49.55 ± 2.95 |
| **10** | 56.2 ± 4.2 | 1.885 ± 0.10 | **82.9 ± 1.7** | **27.95 ± 5.10** |
| **16** | 64.3 ± 3.8 | 0.990 ± 0.06 | 59.0 ± 1.5 | 52.50 ± 4.51 |
| **17** | 47.2 ± 1.0 | 2.067 ± 0.08 | **82.3 ± 2.1** | **15.24 ± 4.58** |
| **20** | 62.6 ± 1.7 | 1.038 ± 0.08 | 69.0 ± 3.5 | 37.81 ± 3.28 |
| **21** | 58.3 ± 2.4 | 1.288 ± 0.05 | 70.6 ± 1.4 | 27.37 ± 2.35 |

## 3. Conclusions

With increasing concerns about upcoming waves of new SARS-CoV-2 infections, the need for an effective and safe therapy against coronaviral diseases is growing. A promising strategy involves M^pro^ inhibition and this approach can lead to novel, broad spectrum anticoronaviral drugs.^13^ Recently, repurposing efforts enabled identification of ebselen as a potential drug against COVID-19, due to its action as a potent inhibitor of the SARS-CoV-2 main protease.^7^ Ebselen has a pleiotropic mode of action that is a result of its reactivity towards cysteine residues affecting many biological pathways^24,26^ but on the other hand is a well-known substance, the efficacy and safety in humans of which has been evaluated in various studies.^36–38^ We utilized a collection of ebselen derivatives to find potent SARS-CoV-2 PL^pro^ and M^pro^ inhibitors. First, we screened a library of organoselenium compounds. Next, for selected compounds, we determined IC_50_ values. The most potent PL^pro^ inhibitor, 2-(3-hydroxypyridin-2-yl)-1,2-benzoselenazol-3-one, displayed the highest potency in the screening assay and the lowest IC_50_ value (0.578 μM). IC_50_ determination enabled identification of two more compounds with inhibitory properties similar to ebselen. These compounds were 2,4- and 2,5-disubstituted derivatives of ebselen that displayed lower potency during screening, but also slightly lower IC_50_ parameters than for the reference inhibitor. A similar analysis for the M^pro^ enabled identification of four compounds displaying higher potency during screening and a lower IC_50_ parameter. Two of them had a monosubstituted phenyl ring at the *para* and *ortho* positions, and two were disubstituted ebselen derivatives. The best inhibitor with an IC_50_ value approx. 2 times lower than for ebselen was 2-(5-chloro-2-fluorophenyl)-1,2-benzoselenazol-3-one. In this work, we showed that ebselen derivatives with substitutions and other modifications within the phenyl ring generally possess good inhibitory properties against both the proteases encoded by the novel coronavirus. These compounds constitute a promising platform for novel therapeutics and we believe that the results obtained can be used to facilitate efforts towards new anticoronaviral drugs to be used for the treatment of COVID-19.

## 4. Materials and methods

### 4.1. SARS-CoV-2 PL^pro^preparation

SARS-CoV-2 PL^pro^ was prepared as described.^22^ In brief, *pGEX6P-1-SARS-CoV-2PLpro* was transformed into BL21 (DE3) codon plus *E. coli* cells and induced with 0.1 mM IPTG and 0.1 mM ZnSO_4_ at 18°C overnight. GST-fusion SARS-CoV-2 PL^pro^ was purified using standard protocol. The fusion protein was cleaved using GST-PreScission protease at 4°C overnight followed with desalting and passing through fresh glutathione beads to remove cleaved GST and GST-PreScission protease. The sample was further purified using Superdex 200 pg size-exclusion columns (GE) equilibrated with 20 mM Tris-Cl pH 8.0, 40 mM NaCl and 2 mM DTT. The peak fractions were pooled and concentrated to ~10 mg/ml and snap frozen in liquid nitrogen for later use.

### 4.2. SARS-CoV-2 M^pro^ preparation

SARS-CoV-2 M^pro^ was recombinantly produced as described.^13^ Briefly, the gene of the M^pro^ was cloned into the PGEX-6p-1 vector, which has a Nsp4-Nsp5 and a PreScission cleavage site at the N- and C-termini to generate the authentic target protein, respectively. The gene of the target protein was expressed in the *E. coli* of the BL21-Gold (DE3) (Novagen) strain. The recombinantly produced M^pro^ was purified by employing HisTrap FF (GE Healthcare) and ion-exchange chromatography (Q FF, GE Healthcare), respectively. Finally, the target protein with high purity was subjected to a buffer exchange (20 mM Tris, 150 mM NaCl, 1 mM EDTA, 1 mM DTT, pH 7.8) for further experiments.

### 4.3. Synthesis of ebselen derivatives

The synthesis of biologically active organoselenium compounds is of current interest to many research teams around the world.^39–41^ Ebselen and other benzisoselenazol-3(2*H*)-ones have been previously prepared by several ways.^42,43^ In this work, we successfully synthesized ebselen and benzisoselenazol-3(2*H*)-ones **1–23** functionalized at the *N*-2 position of the aryl ring, using a four-step procedure previously described in literature, starting with anthranilic acid and elemental selenium.^35,44–46^ The diazotation of previously protonated anthranilic acid, selenentylation with freshly prepared disodium diselenide gave 2,2’-dicarboxydiphenyl diselenide which was isolated before the reaction with thionyl chloride (SOCl_2_) to form 2-(chloroseleno)benzoyl chloride, which is a key substrate in the synthesis of the benzisoselenazol-3(2*H*)-one unit. Tandem selenenylation/acylation reaction of appropriate anilines used in excess or in stoichiometric amounts in the presence of anhydrous triethylamine base in anhydrous MeCN or DCM provided ebselen and final products **1–23**. As a key aniline reagents we used 2-amino-3-hydroxy-pyridine, aniline and its mono- and disubstituted derivatives. The aniline substituents used in different positions of the benzene ring were Me, Ac, NHAc, NO_2_, CF_3_, OH, OMe and F, Cl, Br and I halides. Ebselen and compounds **1–23**, of which eight are new, were obtained. Purity of all new compounds was characterized.

### 4.4. Inhibitor screening

Evaluation of the compound library for inhibitors of SARS-CoV-2 PL^pro^ and SARS-CoV-2 M^pro^ was carried out in Corning 96-wells plates. For PL^pro^, 1 μL of each compound in DMSO solution was added to the wells. Next, 79 μL of enzyme preincubated for 10 min at 37°C in assay buffer (50 mM Tris, 5 mM NaCl, 0.075% BSA, pH 7.5) was added to each well. The enzyme was incubated with the compounds at 37°C for 30 min. Next, 20 μL Ac-LRGG-ACC substrate in assay buffer was added to the wells. Final concentrations were: 100 nM enzyme, 10 μM substrate and 1 μM tested compounds. In the assay for M^pro^, 1 μL of each compound in DMSO solution was added to the wells. Next, 79 μL of enzyme in assay buffer (50 mM Tris, 1 mM EDTA, pH 7.3)^47^ was added to each well and the plate was incubated at room temperature for 2 min. Next, 20 μL of QS1 substrate in assay buffer was added to the wells. Final concentrations were: 100 nM enzyme, 50 μM substrate and 100 nM tested compounds. Measurements were carried out at 37°C using a Molecular Devices Spectramax Gemini XPS spectrofluorometer. ACC fluorophore release was monitored for 30 min (λ_ex_=355 nm, λ_em_=460 nm). For the further analysis, the linear range of the progress curves was used. Measurements were performed at least in duplicate. Results were presented as mean values of relative enzyme inhibition (%, compared to the control measurement without inhibitor) with standard deviations. During the assays, the DMSO concentration in the wells was <2%.

### 4.5. IC_50_ determination

To determine IC_50_, the relative activity of investigated proteases was assessed in at least 11 different concentrations of selected inhibitors. Initial compound concentrations were found experimentally. Serial dilutions of inhibitors in assay buffers (described above) were prepared on 96-well plates (20 μL of each dilution in wells). For SARS-CoV-2 PL^pro^, 60 μL enzyme preincubated for 10 min at 37°C in assay buffer was added to the wells. The enzyme was incubated with inhibitors for 30 min at 37°C. Next, 20 μL substrate (Ac-LRGG-ACC) in assay buffer was added to the wells. Final concentrations were 100 nM enzyme and 10 μM substrate.

For SARS-CoV-2 M^pro^, 60 μL enzyme was added with no preincubation. The enzyme was incubated with inhibitor for 2 min at room temperature. Next, 20 μL of substrate (QS1) in the assay buffer was added to the wells. Final concentrations were 100 nM for the enzyme and 50 μM for the substrate. Measurements were carried out at 37°C using a Molecular Devices Spectramax Gemini XPS spectrofluorometer. ACC fluorophore release was monitored for 30 min (λ_ex_=355 nm, λ_em_=460 nm). IC_50_ values were determined with GraphPad Prism software using non-linear regression (dose-response - Inhibition equation) and presented as relative enzyme activity vs. inhibitor concentration. Measurements were performed at least in triplicate. Results are presented as mean values with standard deviations. During the assays, the DMSO concentration in wells was <2%.

## Author contributions

M. Z. and M. D. designed the research; M. Z. and W. R. performed the research and collected data; K. O., J. G., M. G. and M. B.-G. synthesized and provided the collection of compounds, M. K.-B., L. Z., X. S. and R. H. provided SARS-CoV-2 M^pro^ enzyme; Z. L., D. N. and S. K. O. provided SARS-CoV-2 PL^pro^ enzyme, M. Z. and W. R. analyzed and interpreted the inhibitory data and M. Z. wrote the manuscript; all authors critically revised the manuscript.

## Acknowledgements

This work was supported by the Medical Research Agency in Poland through its Own Project (grant 2020/ABM/SARS/1) and by the National Science Centre in Poland (2020/01/0/NZ1/00063). The Drag laboratory is supported by the “TEAM/2017-4/32” project, which is conducted within the TEAM programme of the Foundation for Polish Science cofinanced by the European Union under the European Regional Development Fund. Work in the Hilgenfeld laboratory was supported by the SCORE project of the European Union (grant agreement # 101003627), by the German Center for Infection Research (DZIF), and by the Government of Schleswig-Holstein through its Structure and Excellence Fund, as well as by a close partnership between the Possehl Foundation (Lübeck) and the University of Lübeck. L.Z. is supported by a stipend from the German Center for Infection Research (DZIF). We are thankful to Wroclaw University of Science and Technology for support for the Department of Organic and Medicinal Chemistry – K20 (Statute Founds 82013902).

