## Supplemental for "Ebselen derivatives are very potent dual inhibitors of SARS-CoV-2 proteases - PL^pro^ and M^pro^ in in vitro studies"

Content of supporting information:

1. LC-MS analysis of investigated compounds .......... S2-S25

#### Mass spectra and analytical chromatograms of the compounds

The purity and molecular weight of each compound was confirmed with LC-MS system (Waters e2695 Separations Module, 2489 UV/Vis Detector, Acquity QDa MS Detector). Analytical HPLC column: Jupiter 10 mm C4 300 Å (250x4.6 mm). Solvent composition: phase A (water/0.1% HCOOH) and phase B (ACN/0.1% HCOOH); gradient from 5% B to 95% B over a period of 15 min.

##### 2-phenylbenzisoselenazol-3(2H)-one (ebselen)

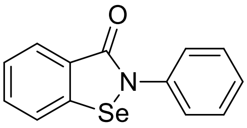


($m/z_{calcd}$ = 275.9923; $m/z_{found}$ = 275.87)

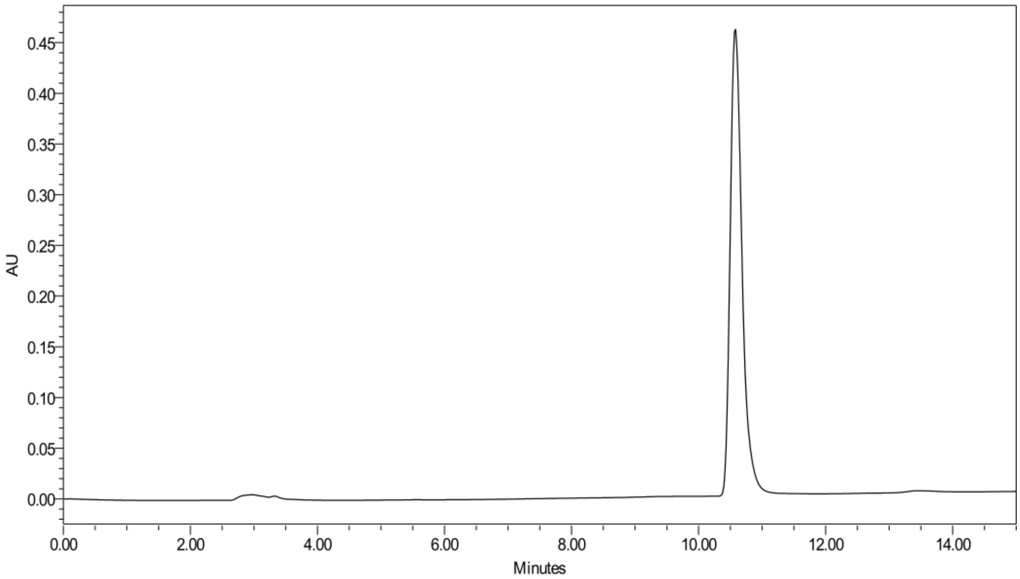


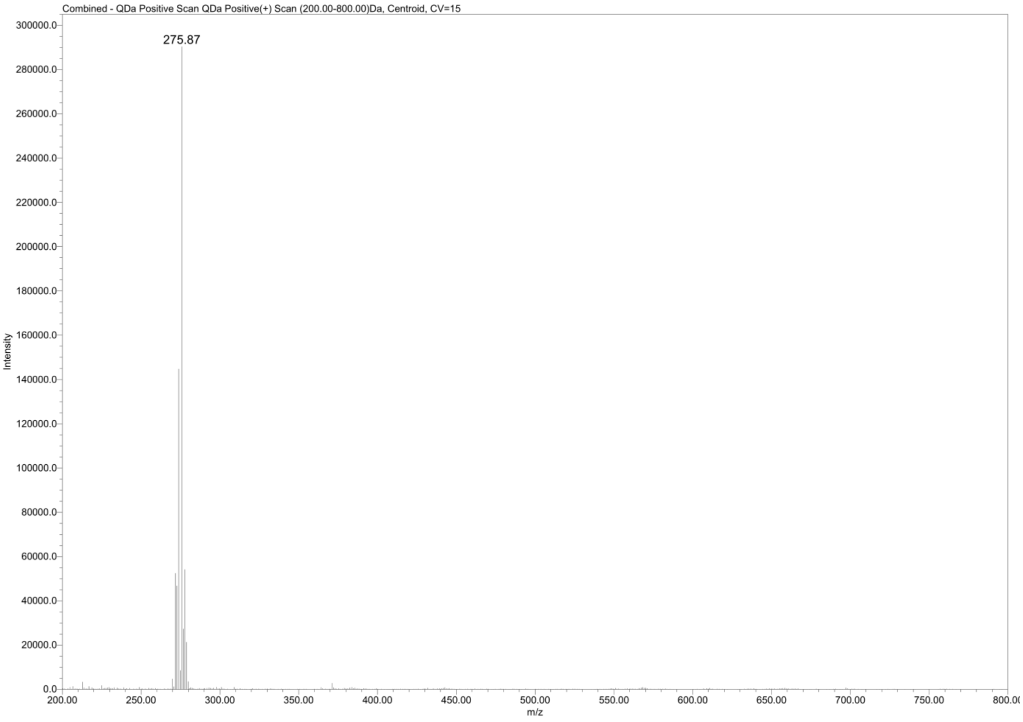

**Compound 1**

**2-(2-fluorophenyl)-benzisoselenazol-3(2H)-one**

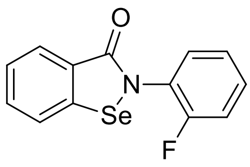


(m/z $_{calcd}$ = 293.9828; m/z $_{found}$ = 293.81)

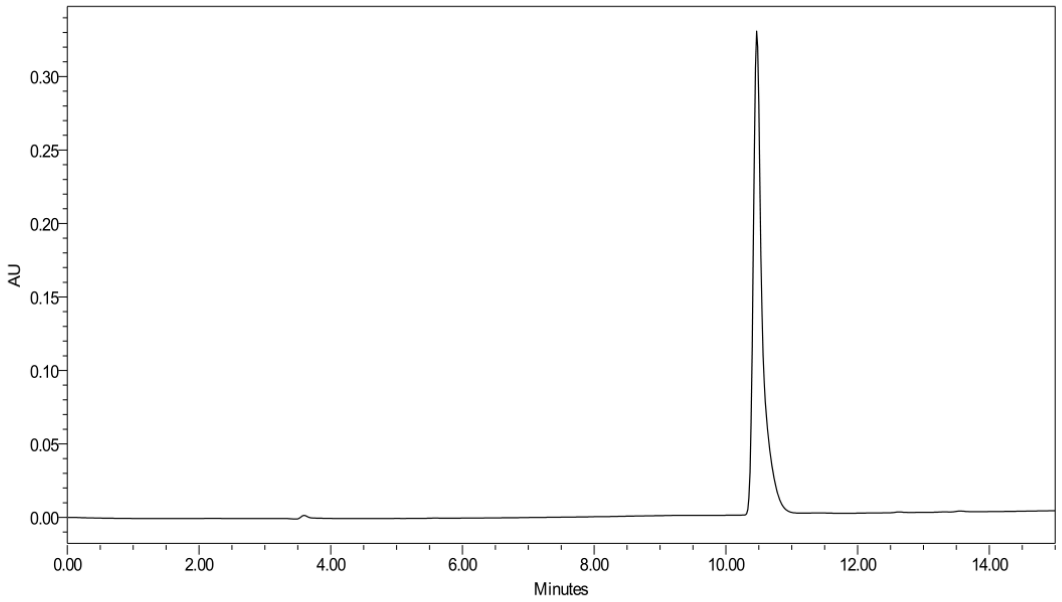


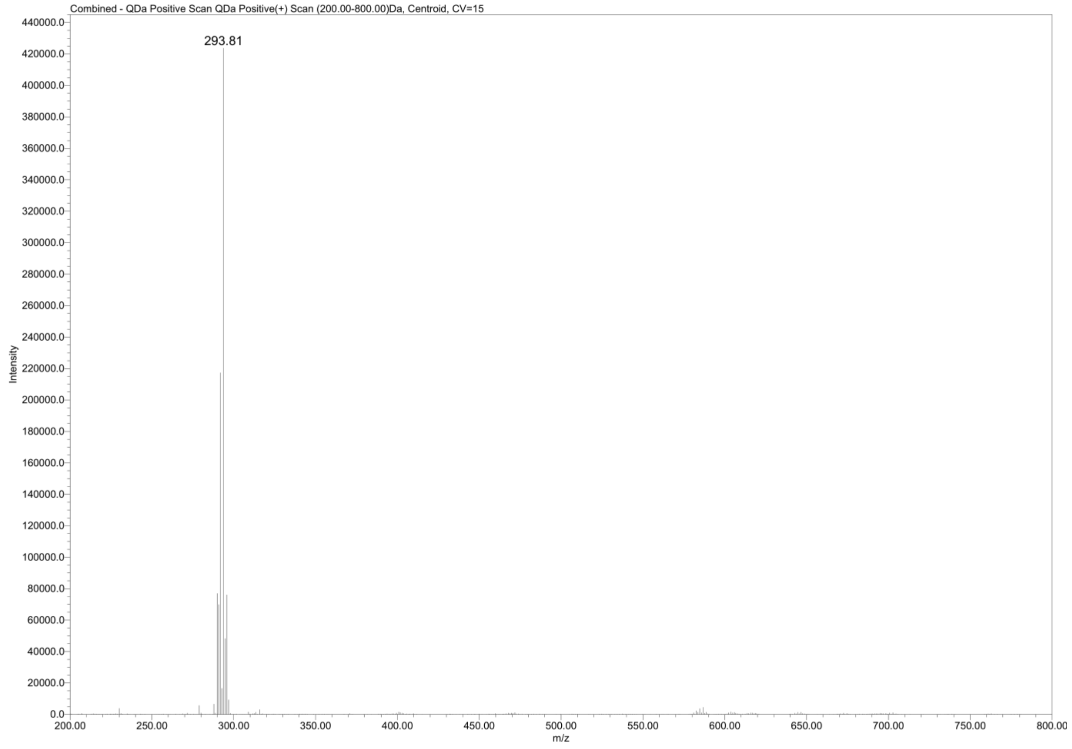

**Compound 2**

**2-(2-chlorophenyl)-benzisoselenazol-3(2H)-one**

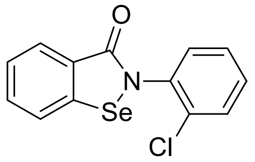


($m/z_{calcd}$ = 309.9533; $m/z_{found}$ = 309.79)

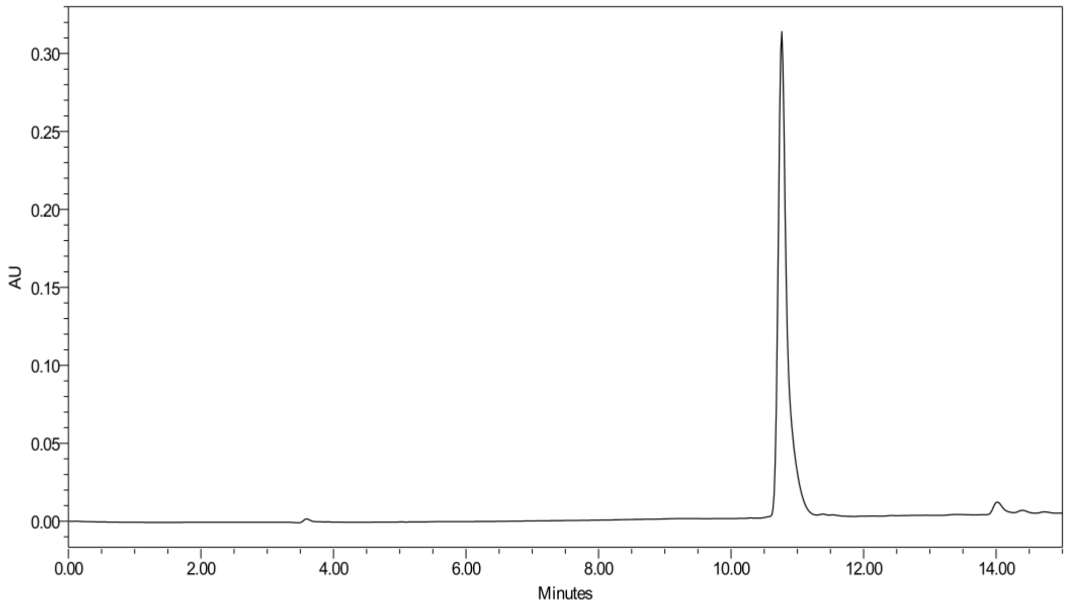


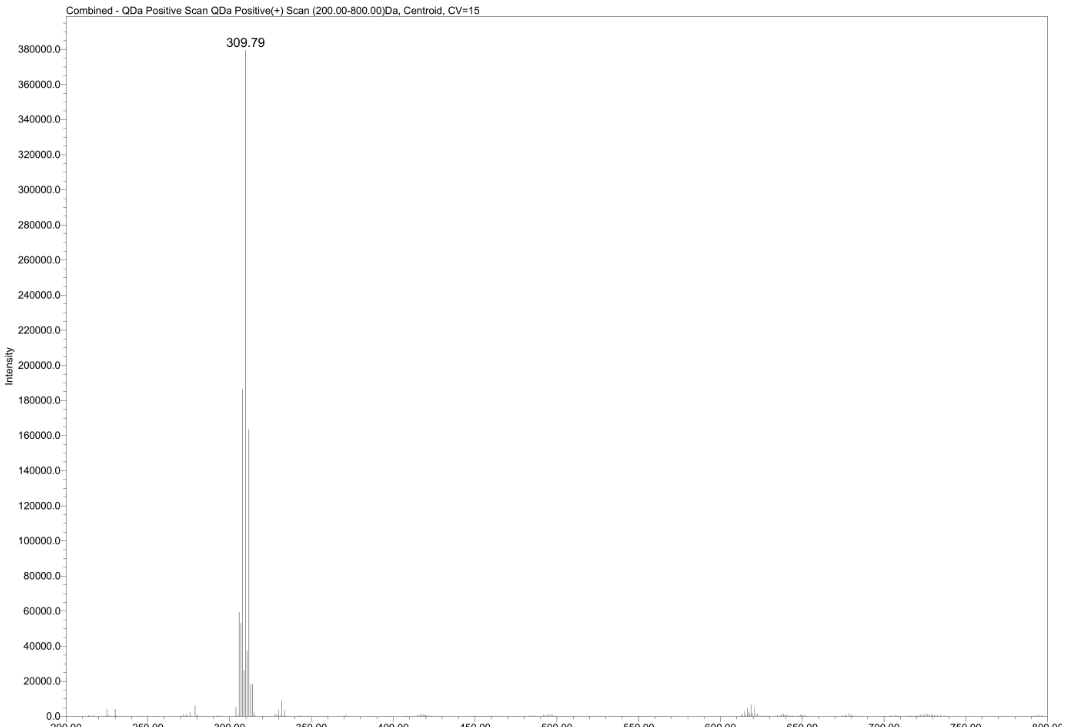

**Compound 3**

**2-(2-bromophenyl)-benzisoselenazol-3(2H)-one**

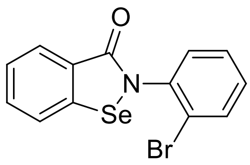


(m/z $_{calcd}$ = 353.9028; m/z $_{found}$ = 353.74)

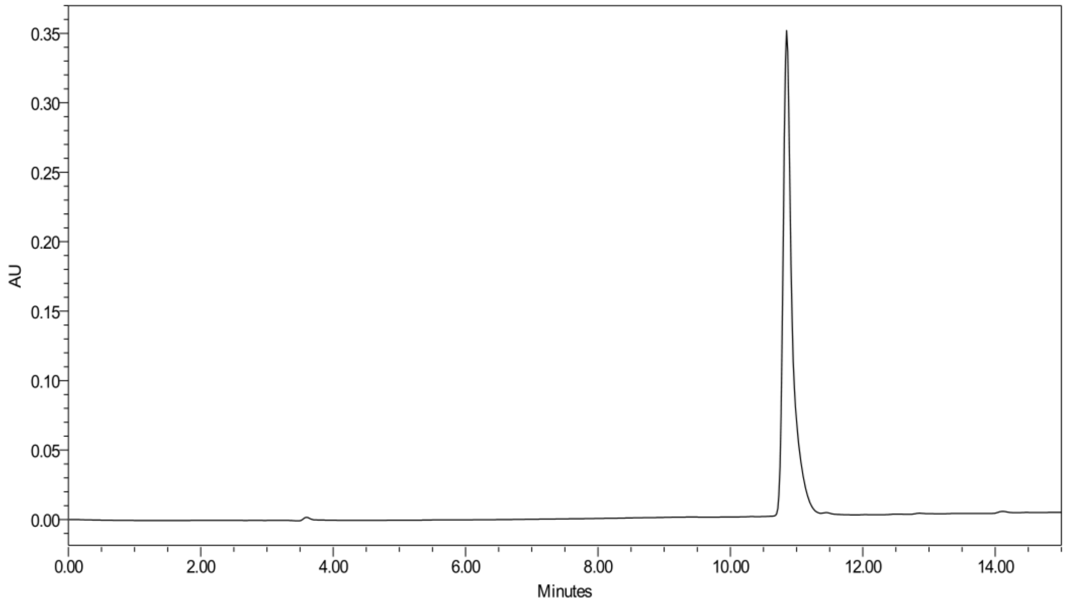


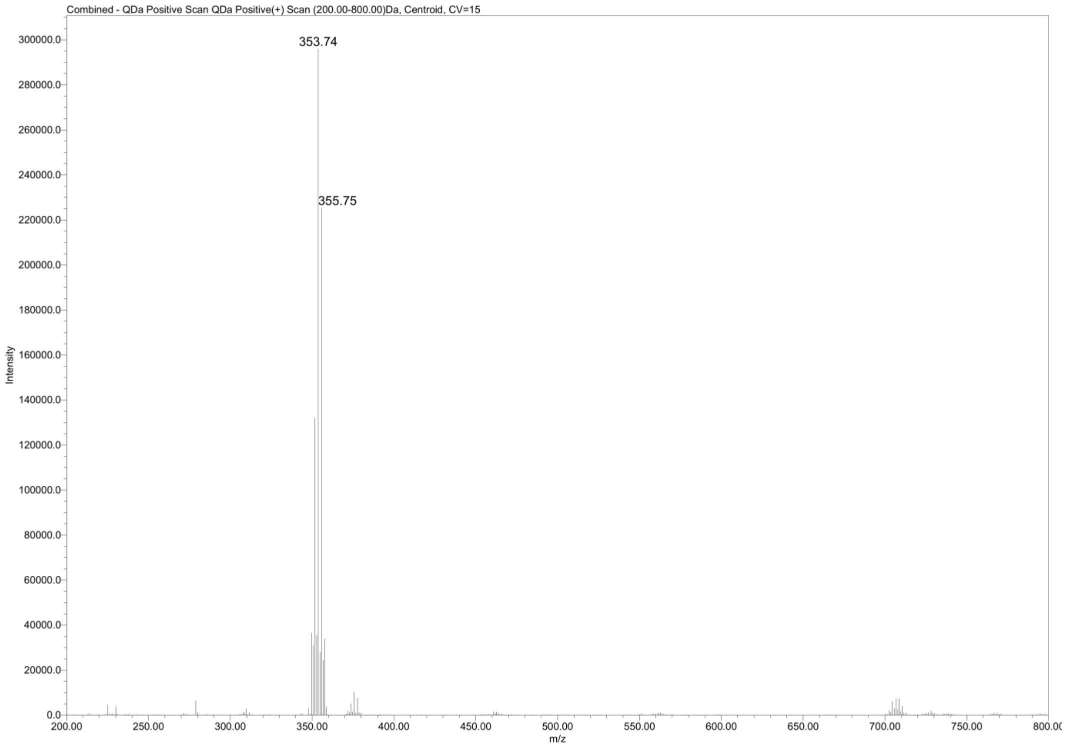

**Compound 4**

**2-(2-methylphenyl)-benzisoselenazol-3(2H)-one**

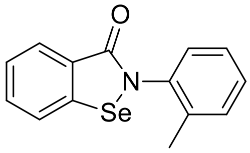


(m/z $_{calcd}$ = 290.0079; m/z $_{found}$ = 289.85)

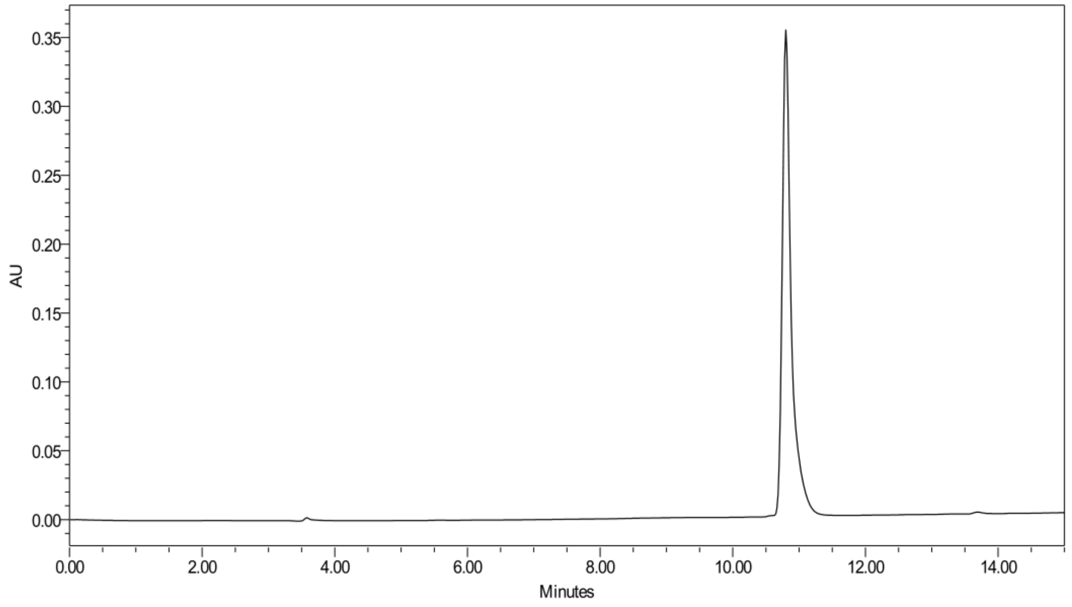


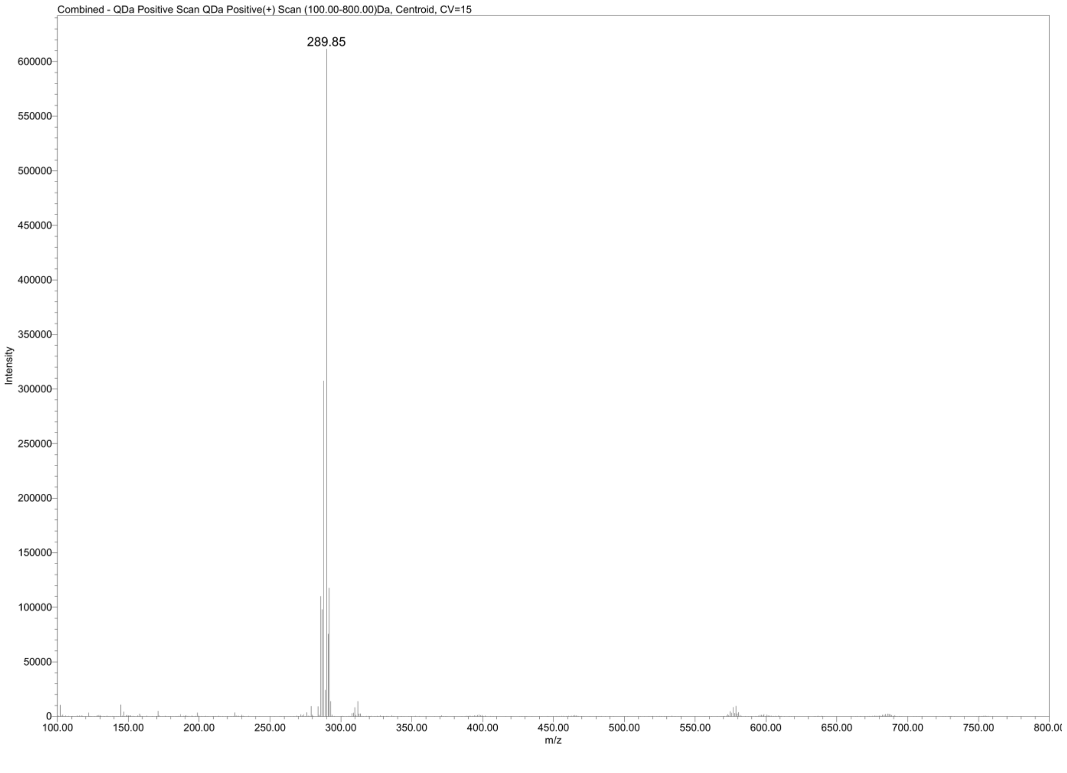

**Compound 5**

**2-(2-(trifluoromethyl)phenyl)-benzisoselenazol-3(2H)-one**

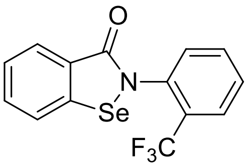


($m/z_{\text{calcd}}$ = 343.9796; $m/z_{\text{found}}$ = 343.84)

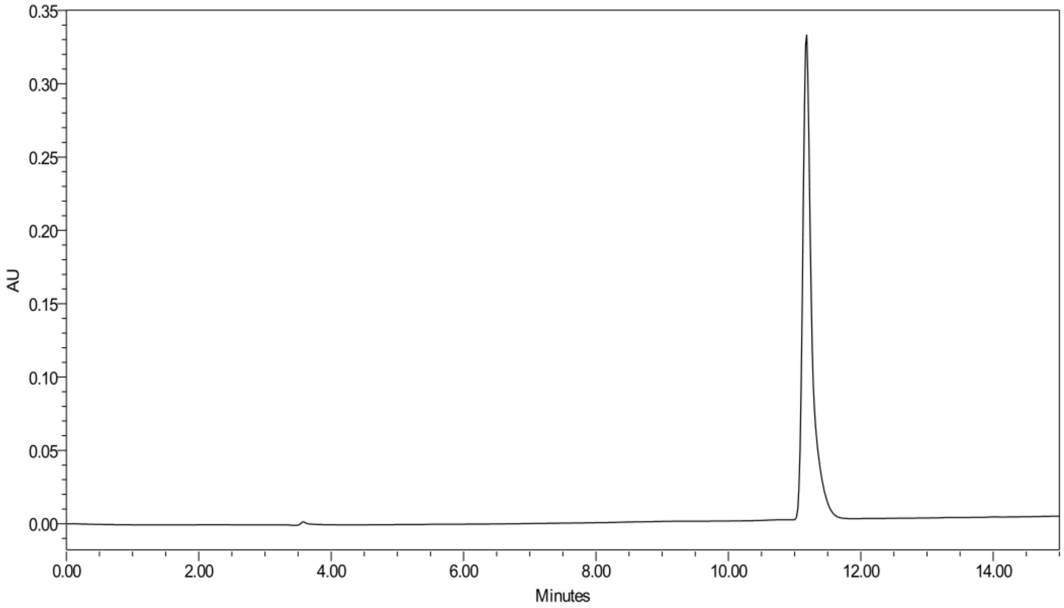


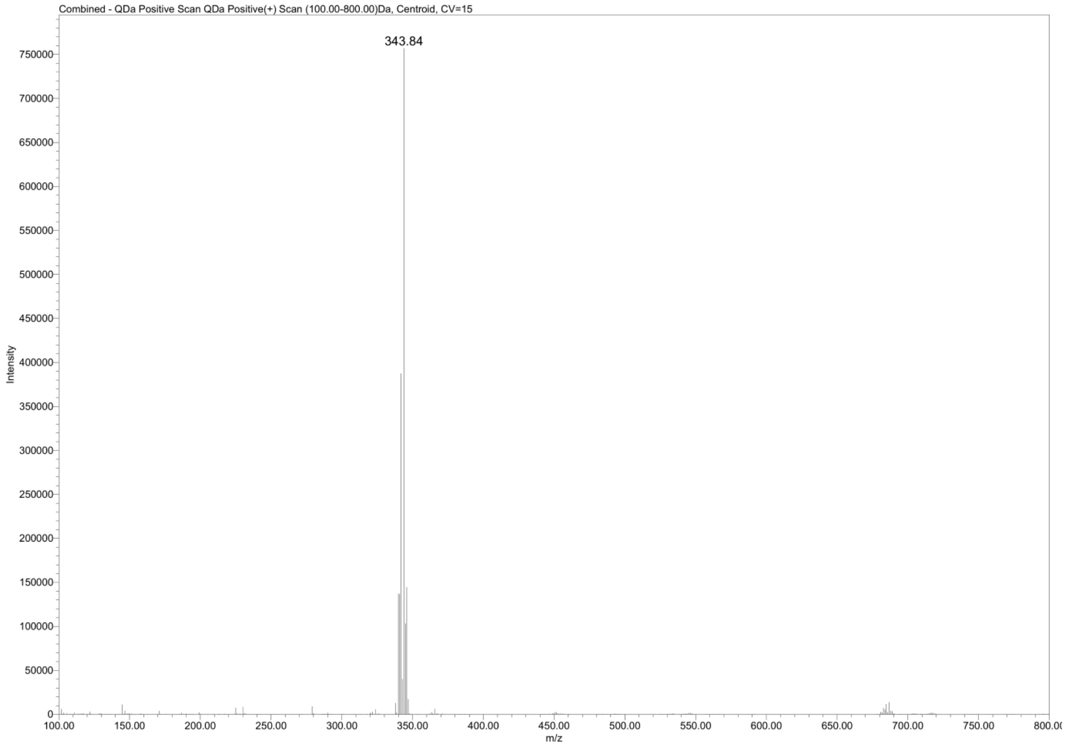

**Compound 6**

**2-(2-nitrophenyl)-benzisoselenazol-3(2H)-one**

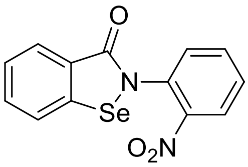

(m/z $_{calcd}$ = 320.9773; m/z $_{found}$ = 320.83)

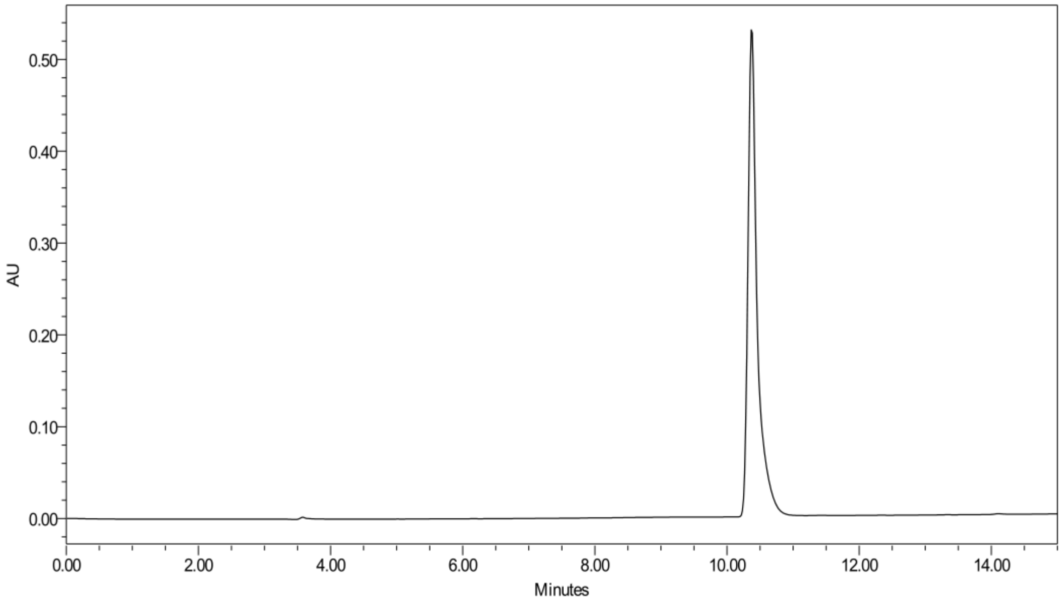


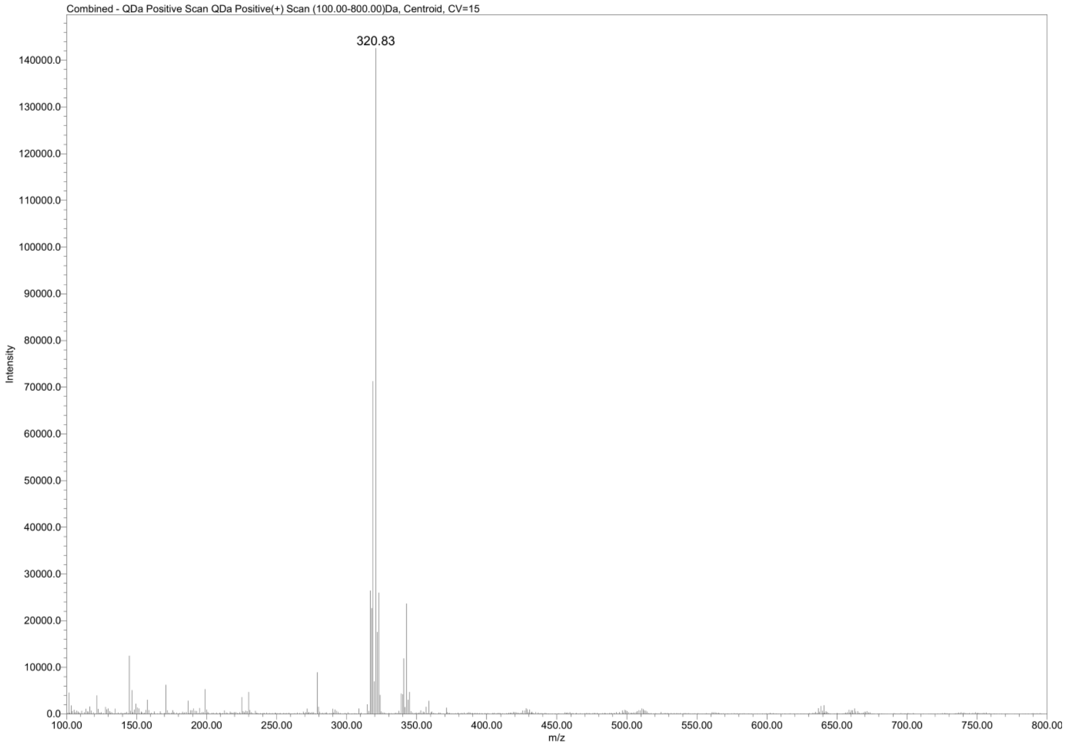

**Compound 7**

**2-(3-hydroxypyridin-2-yl)-1,2-benzoselenazol-3-one**

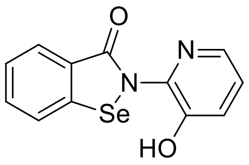


(m/z $_{calcd}$ = 292.9824; m/z $_{found}$ = 292.80)

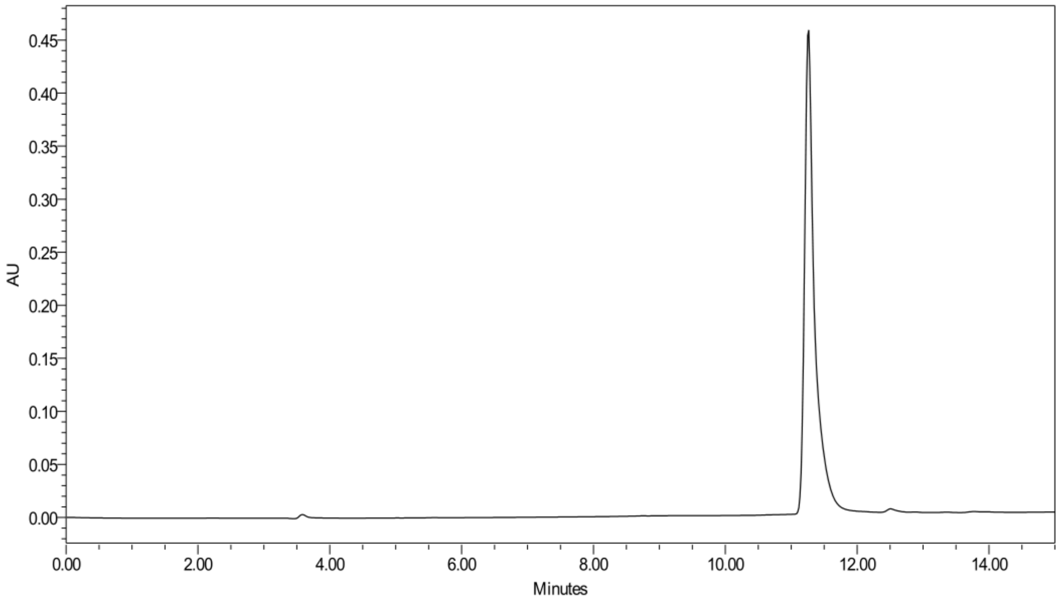


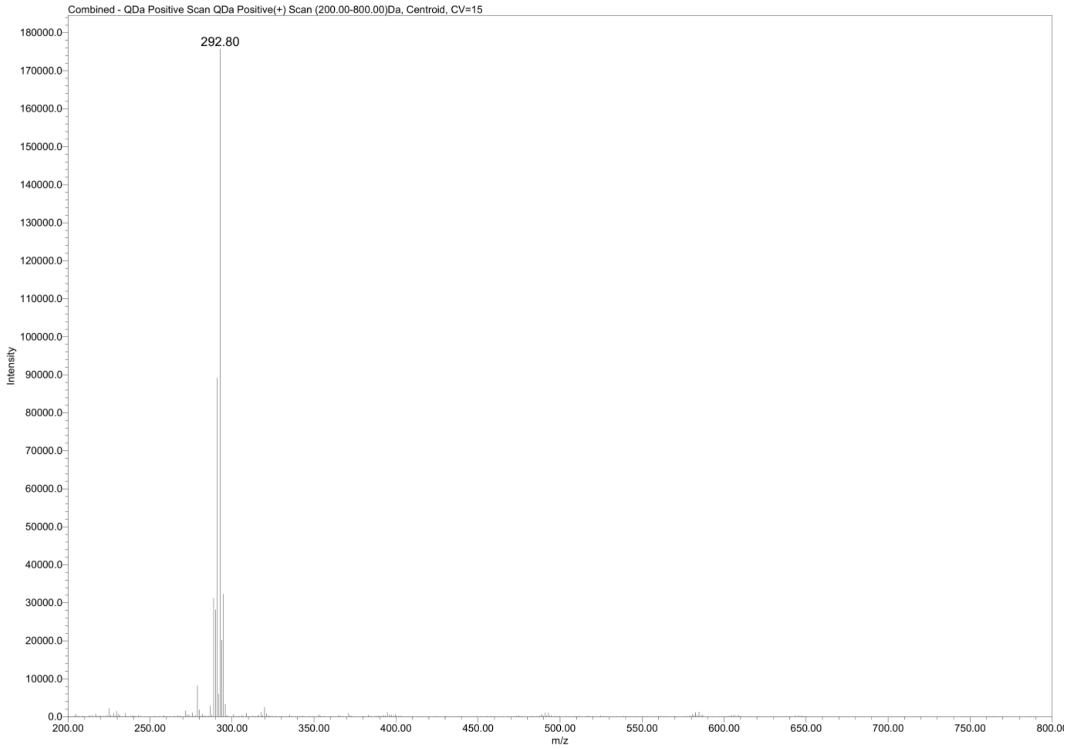

**Compound 8**

**2-(3-methoxyphenyl)-benzisoselenazol-3(2H)-one**

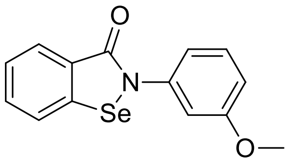


(m/z $_{calcd}$ = 306.0028; m/z $_{found}$ = 305.85)

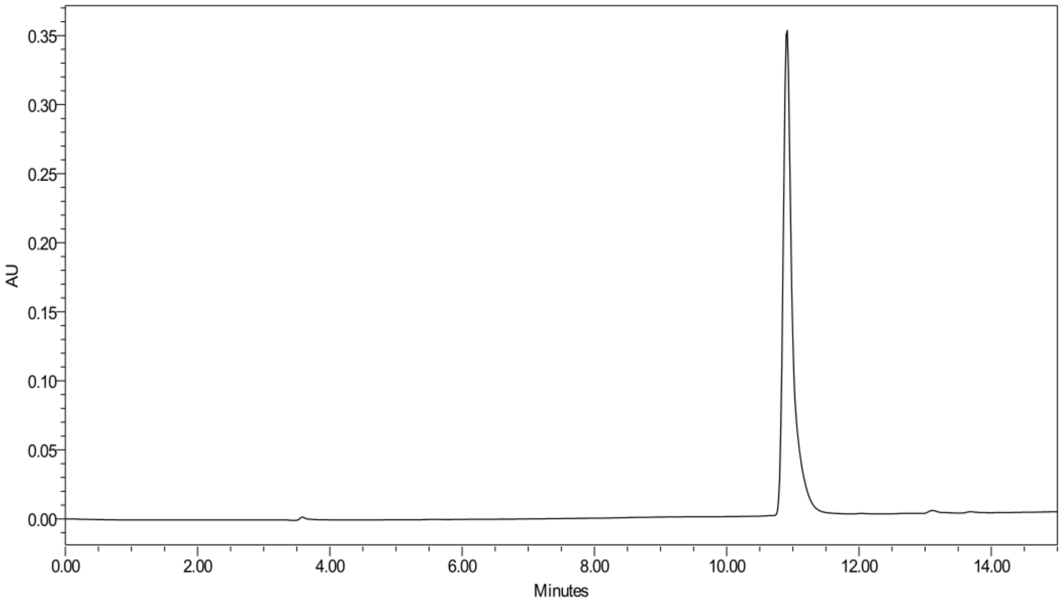


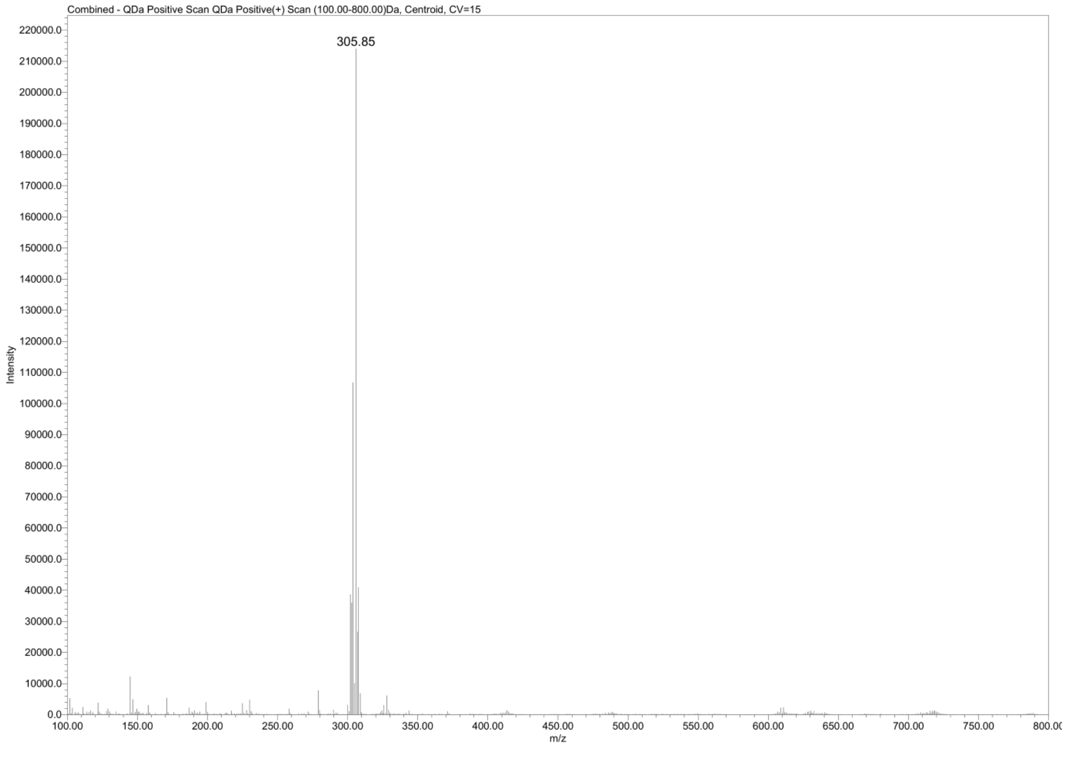

**Compound 9**

**2-(4-(trifluoromethyl)phenyl)-benzisoselenazol-3(2H)-one**

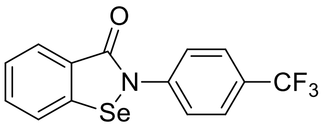


($m/z_{calcd}$ = 343.9796; $m/z_{found}$ = 343.88)

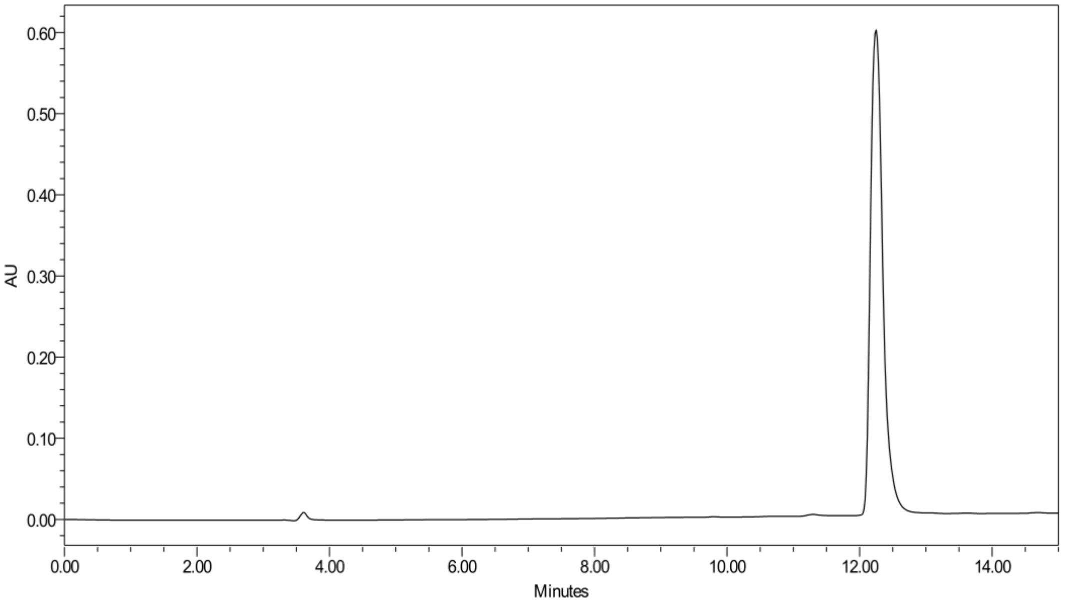


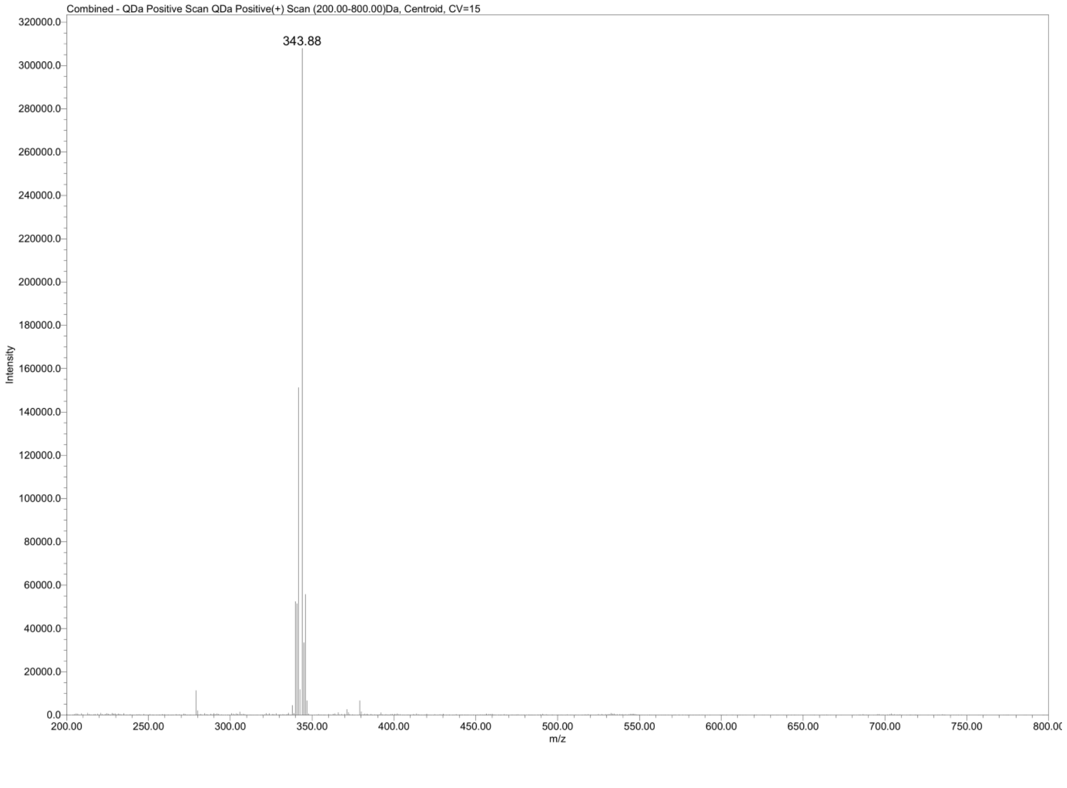

**Compound 10**

**2-(4-nitrophenyl)-benzisoselenazol-3(2H)-one**

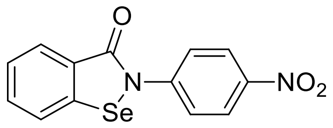


(m/z $_{calcd}$ = 320.9773; m/z $_{found}$ = 320.82)

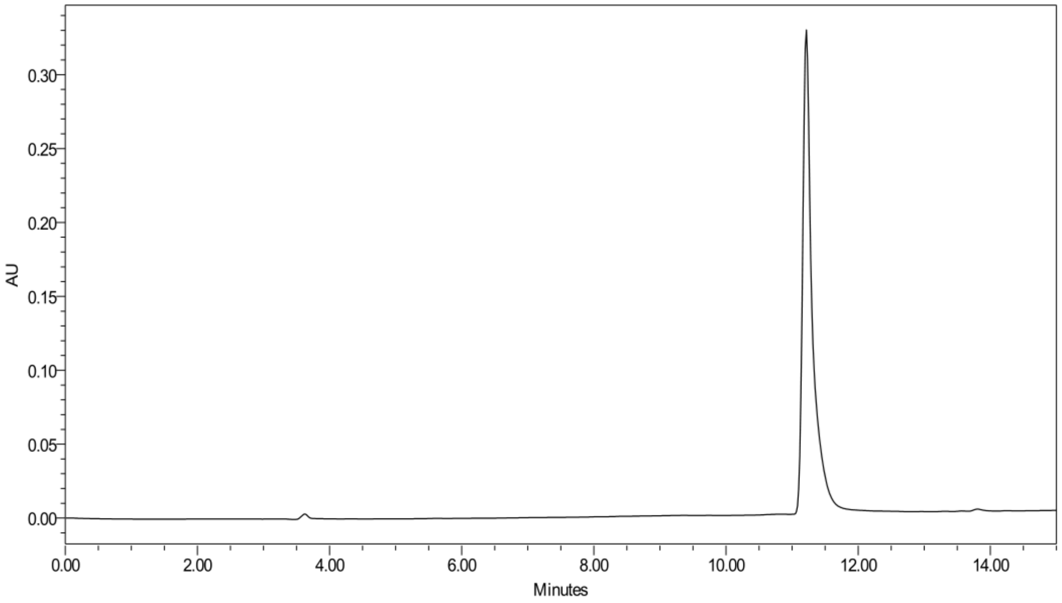


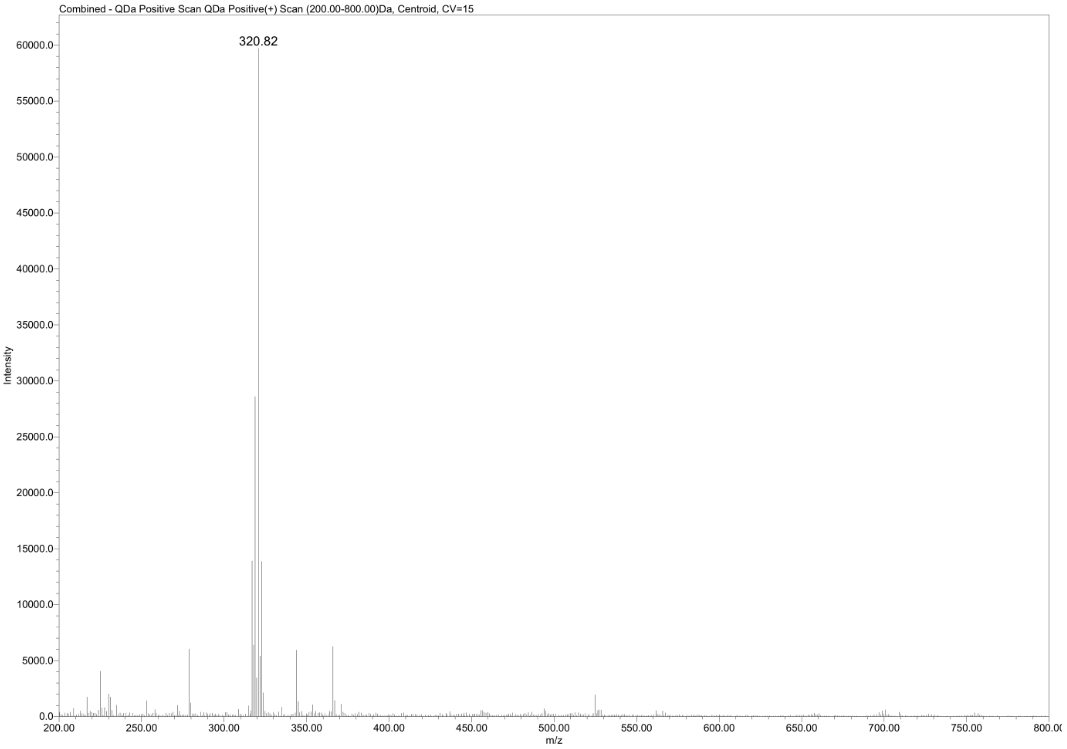

**Compound 11**

**2-(4-iodophenyl)-benzisoselenazol-3(2H)-one**

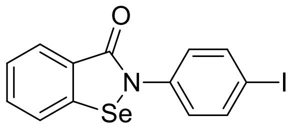


(m/z $_{calcd}$ = 401.8889; m/z $_{found}$ = 401.73)

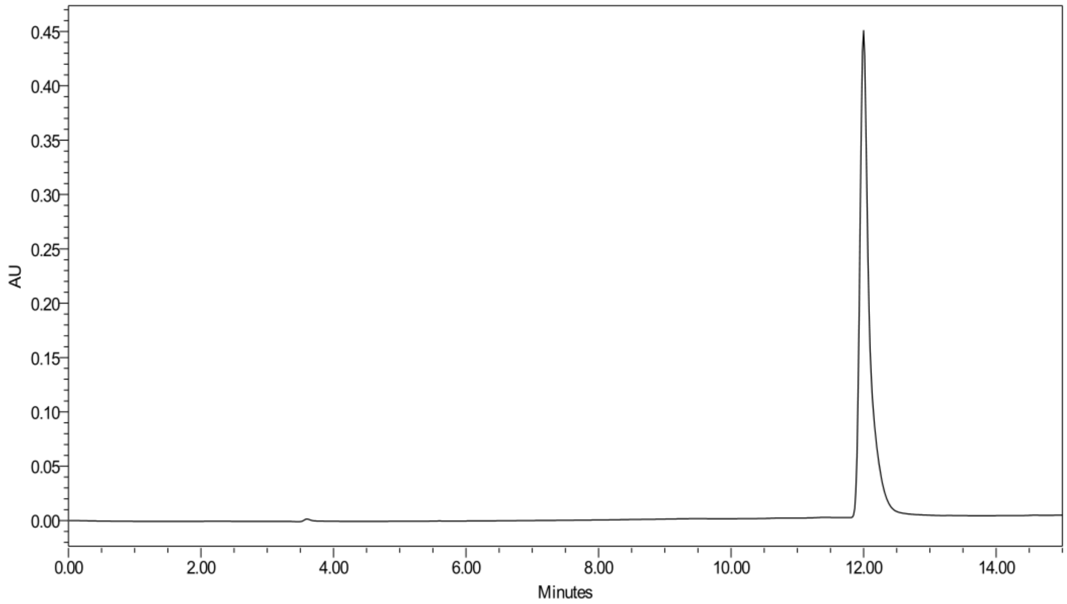


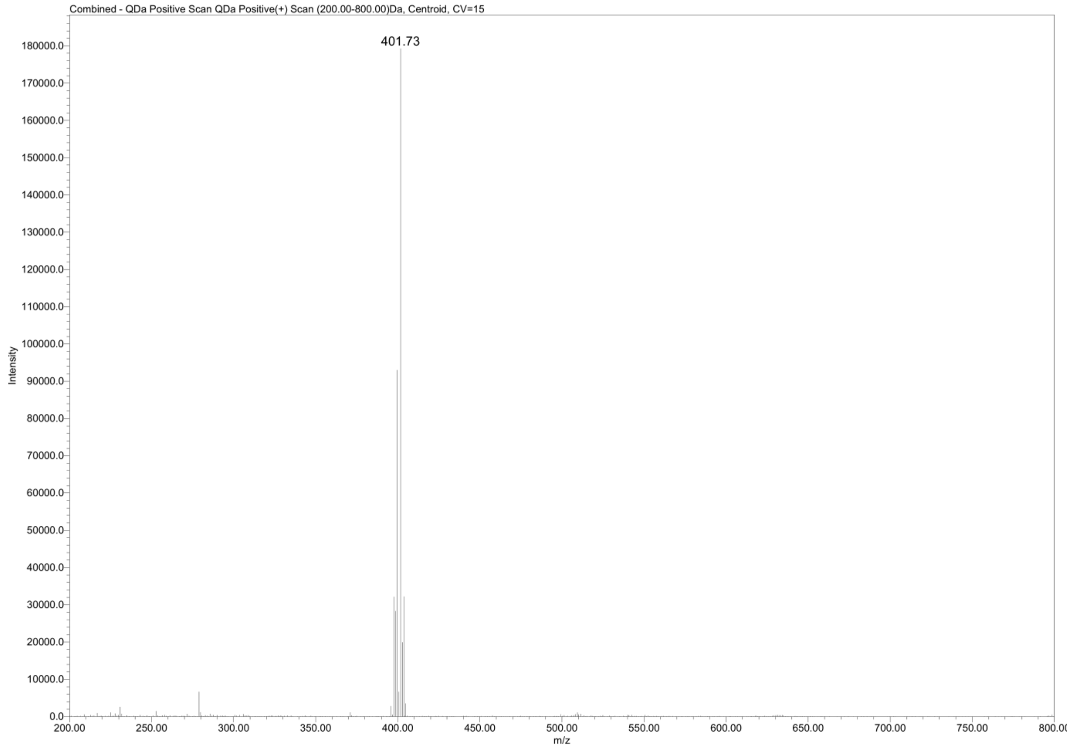

#### Compound 12

#### 2-(4-acetylphenyl)-benzisoselenazol-3(2H)-one

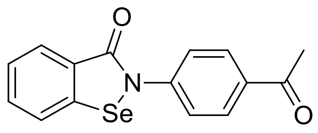


($m/z_{calcd}$ = 318.0028; $m/z_{found}$ = 317.85)

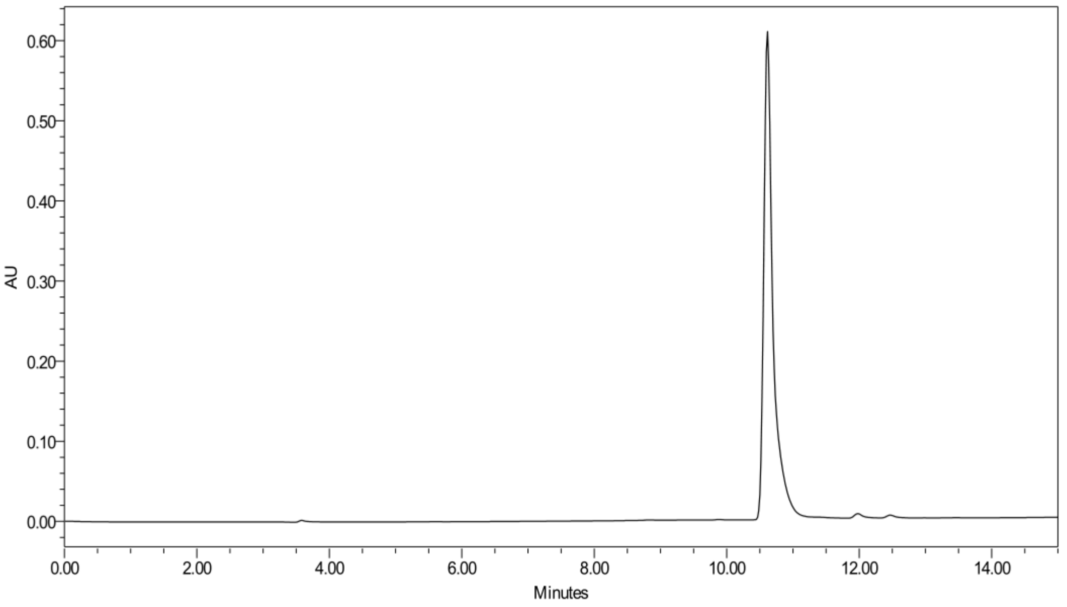


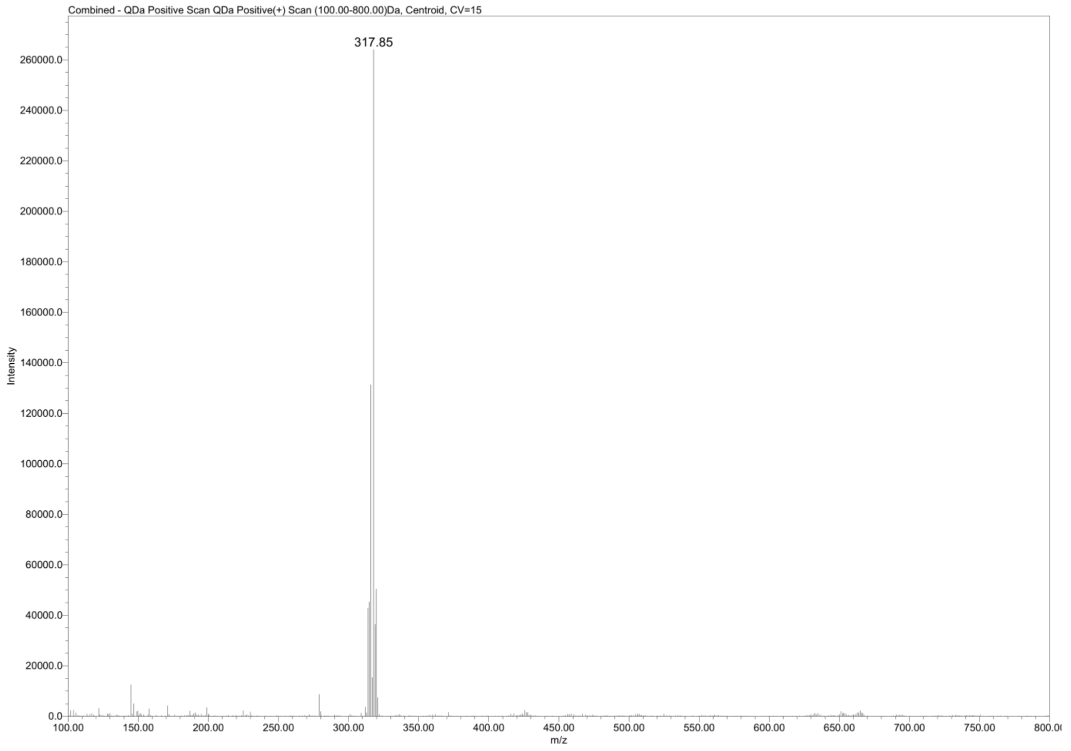

**Compound 13**

**2-(4-acetamidophenyl)-benzisoselenazol-3(2H)-one**

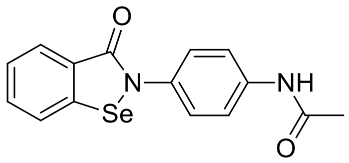


(m/z $_{calcd}$ = 333.0137; m/z $_{found}$ = 332.86)

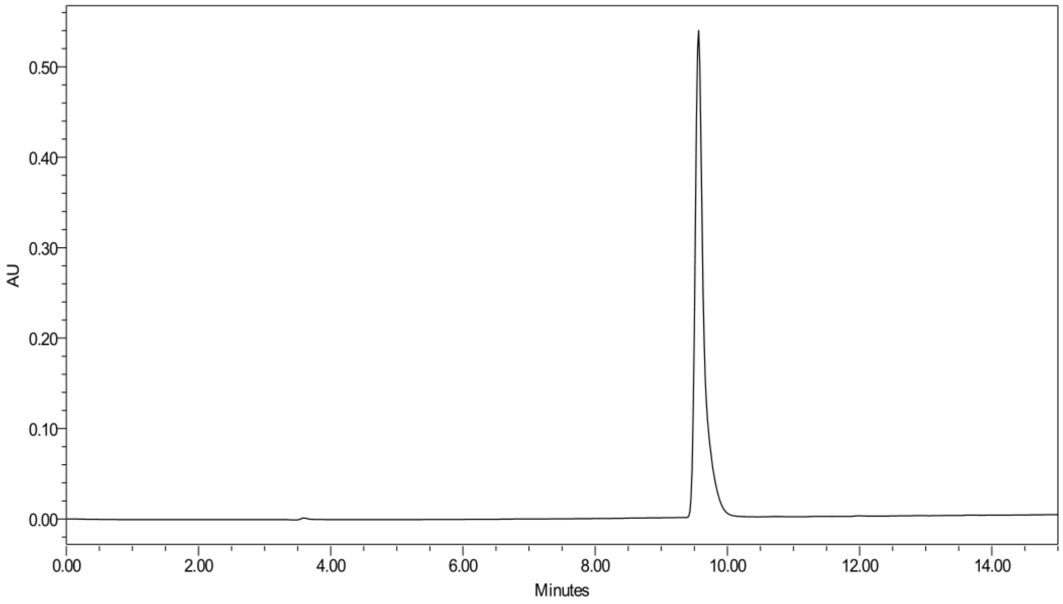


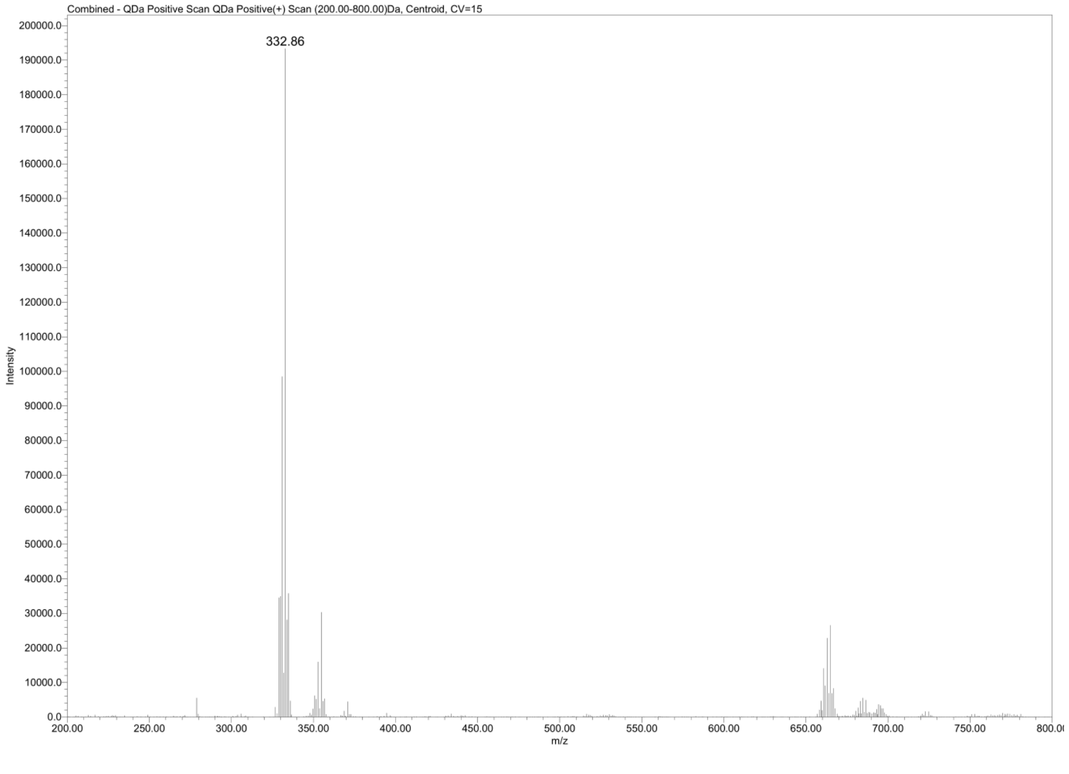

**Compound 14**

**2-(2,4-difluorophenyl)-benzisoselenazol-3(2H)-one**

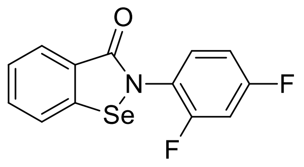


(m/z $_{calcd}$ = 311.9734; m/z $_{found}$ = 311.82)

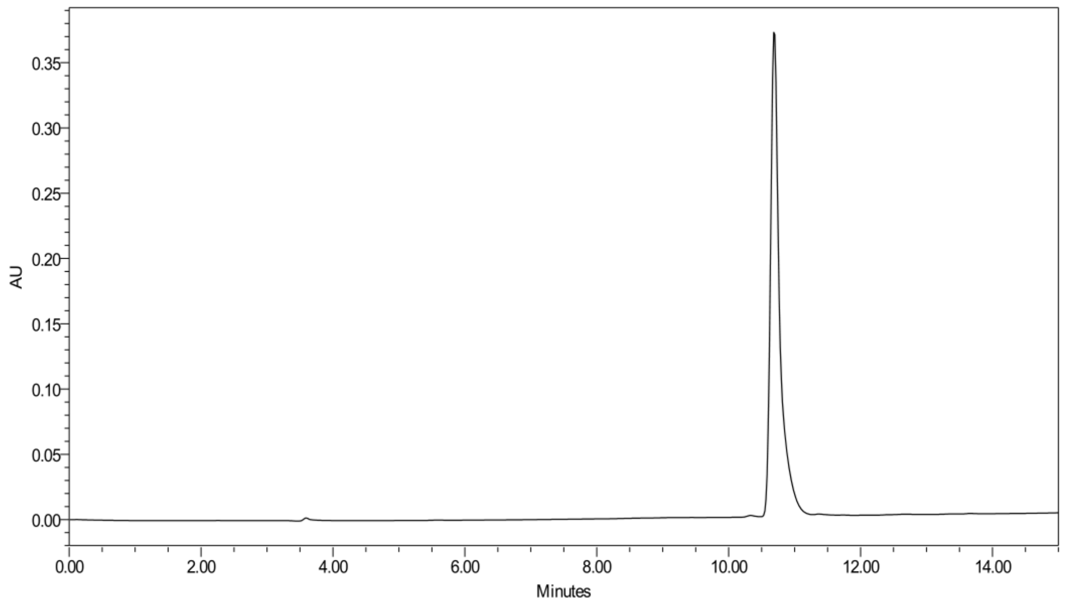


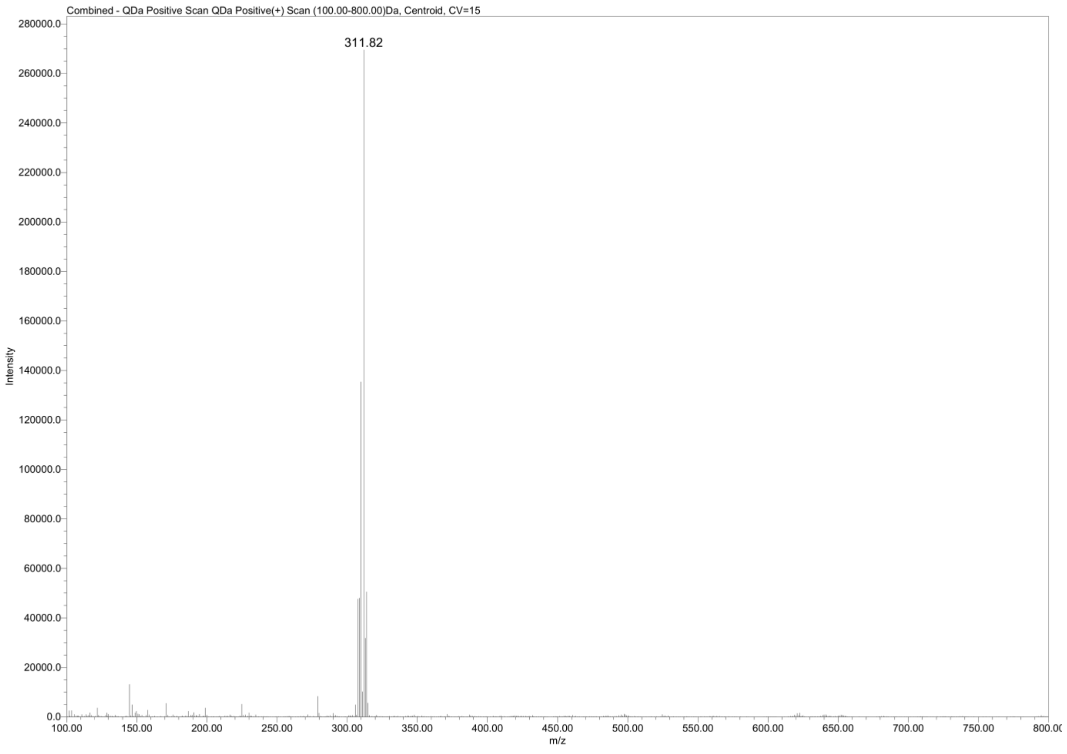

#### Compound 15

#### 2-(4-chloro-2-fluorophenyl)-benzisoselenazol-3(2H)-one

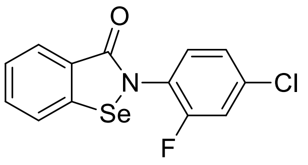


(m/z $_{calcd}$ = 327.9439; m/z $_{found}$ = 327.78)

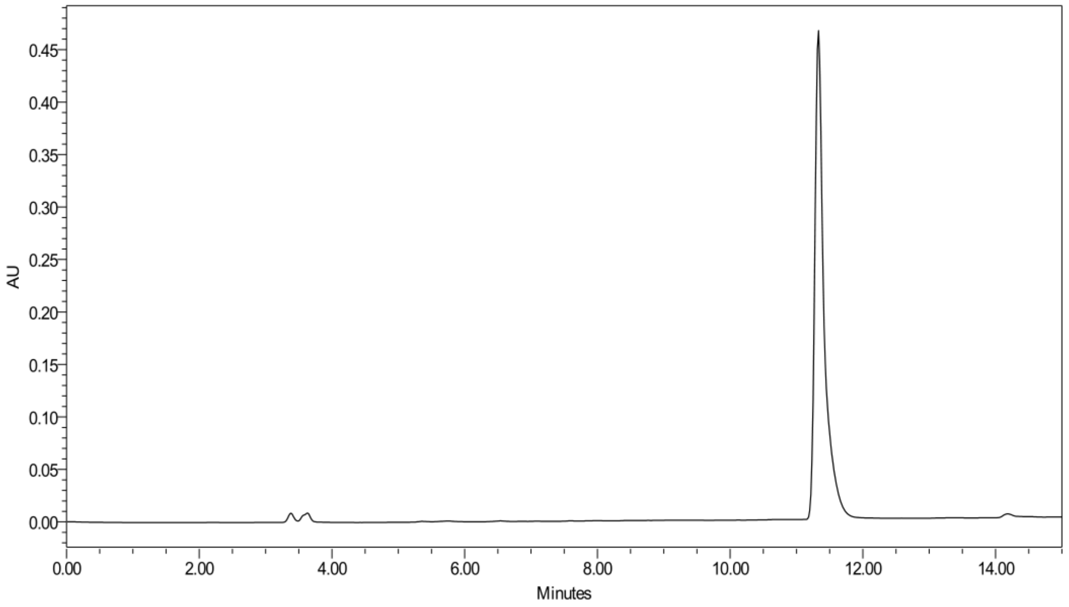


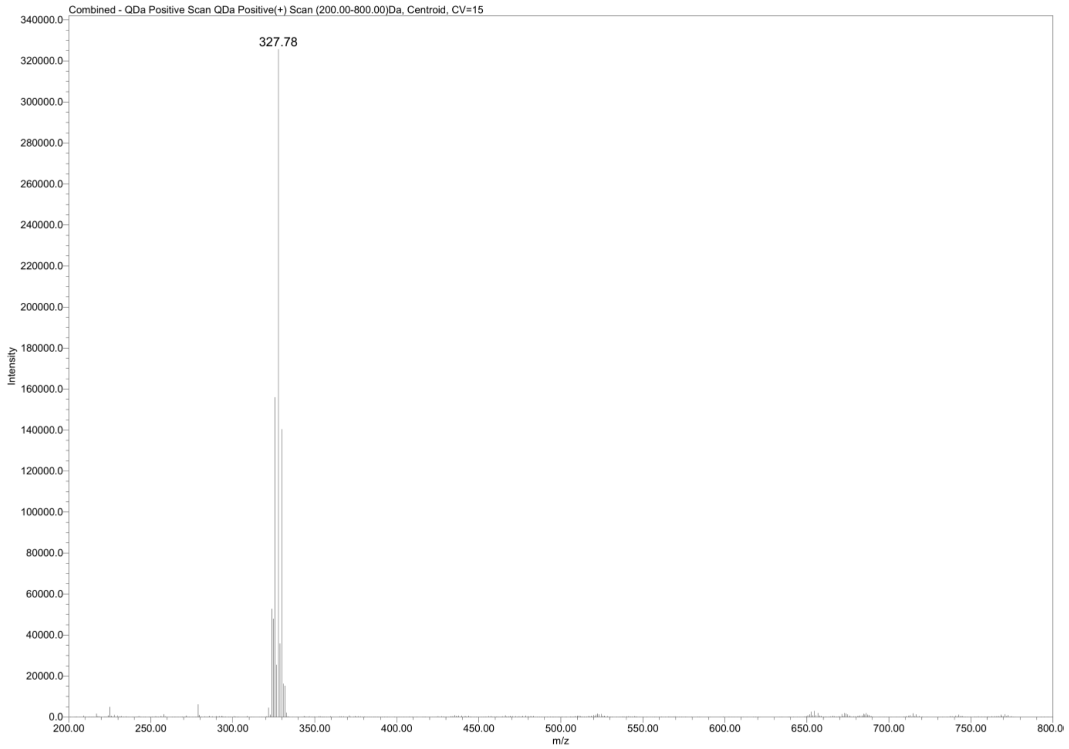

### Compound 16

#### 2-(2,4-dimethoxyphenyl)-benzisoselenazol-3(2H)-one

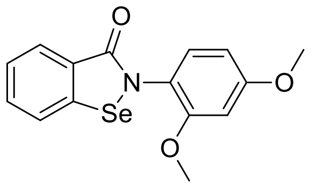


(m/z$_{calcd}$ = 336.0134; m/z$_{found}$ = 335.88)

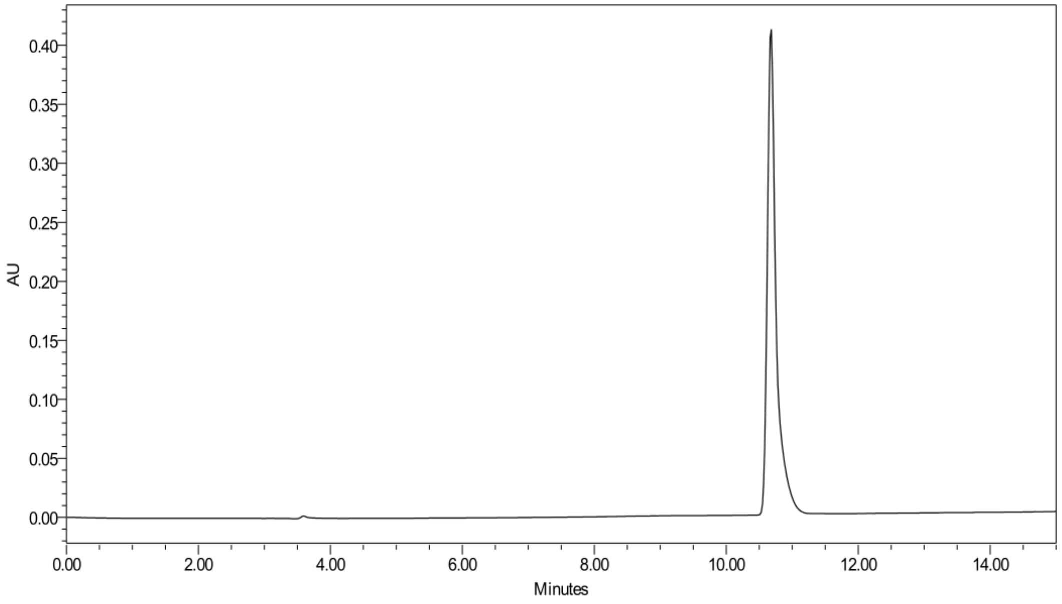


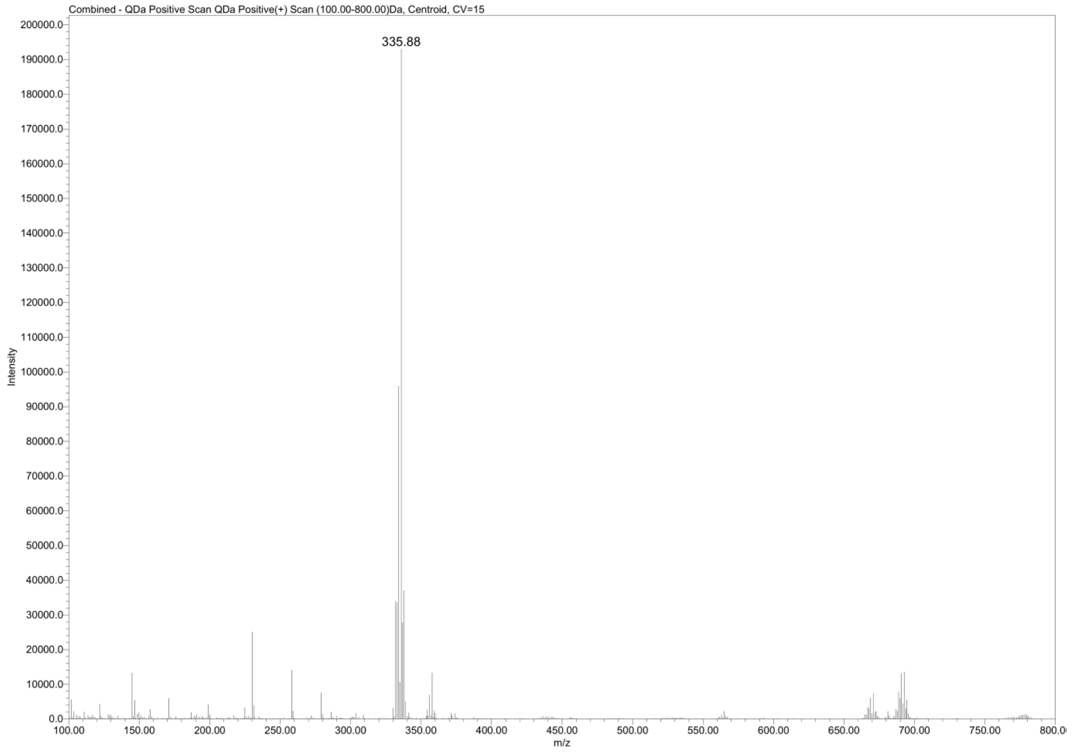

**Compound 17**

**2-(5-chloro-2-fluorophenyl)-benzisoselenazol-3(2H)-one**

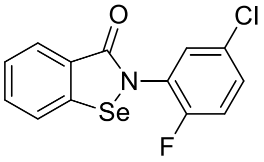


($m/z_{calcd}$ = 327.9439; $m/z_{found}$ = 327.80)

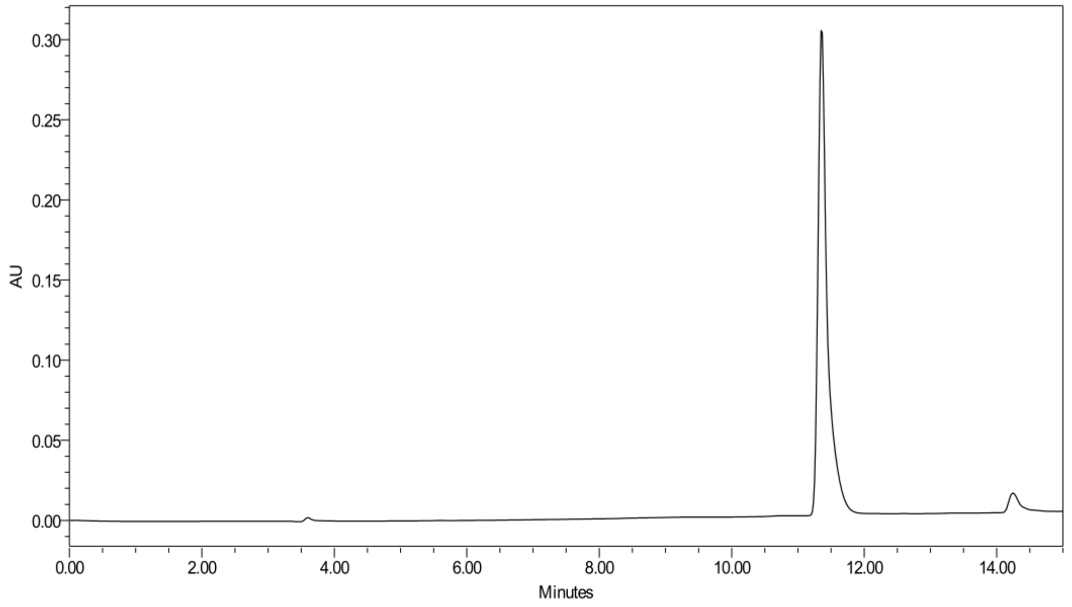


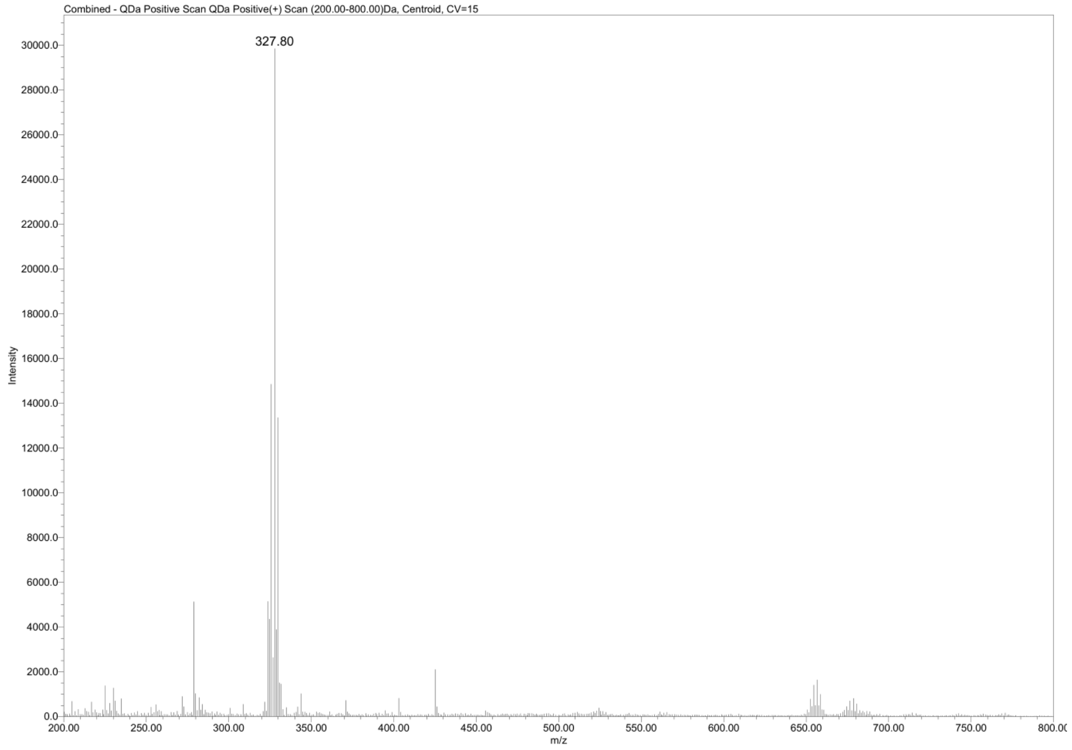

**Compound 18**

**2-(2,5-dichlorophenyl)-benzisoselenazol-3(2H)-one**

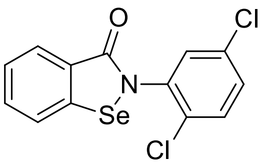


($m/z_{calcd}$ = 343.9143; $m/z_{found}$ = 343.75)

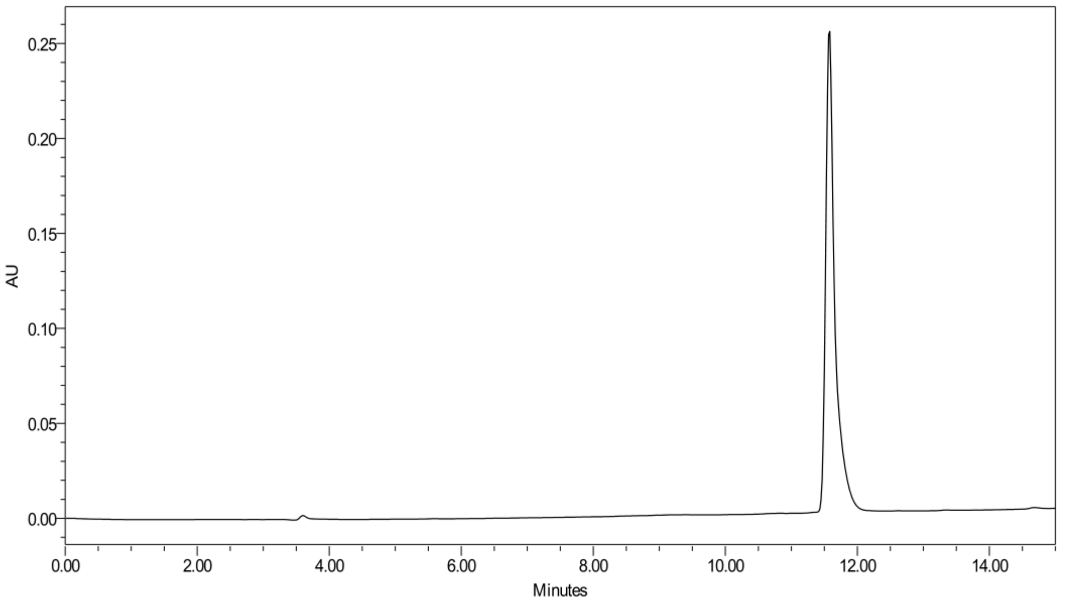


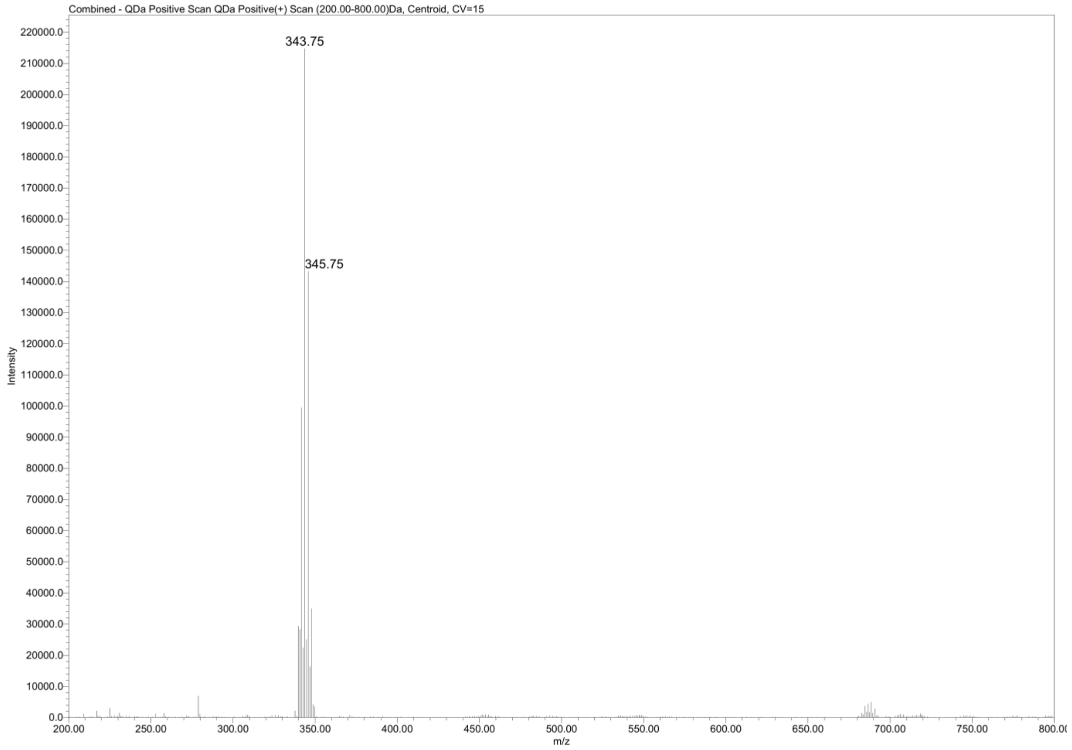

**Compound 19**

**2-(2-chloro-5-methylphenyl)-benzisoselenazol-3(2H)-one**

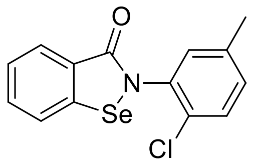


($m/z_{calcd}$ = 323.9689; $m/z_{found}$ = 323.87)

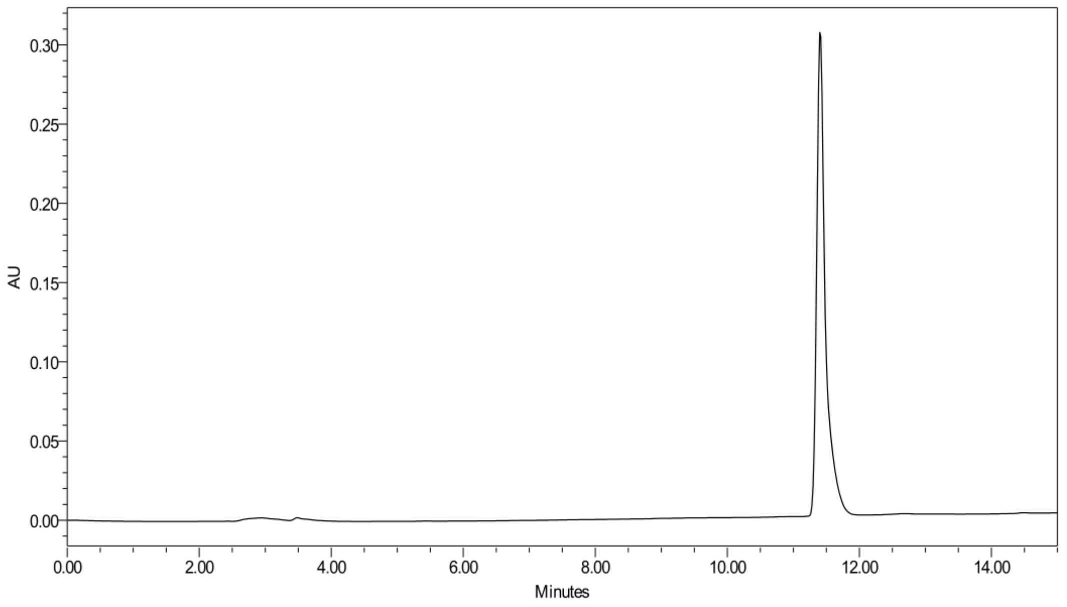


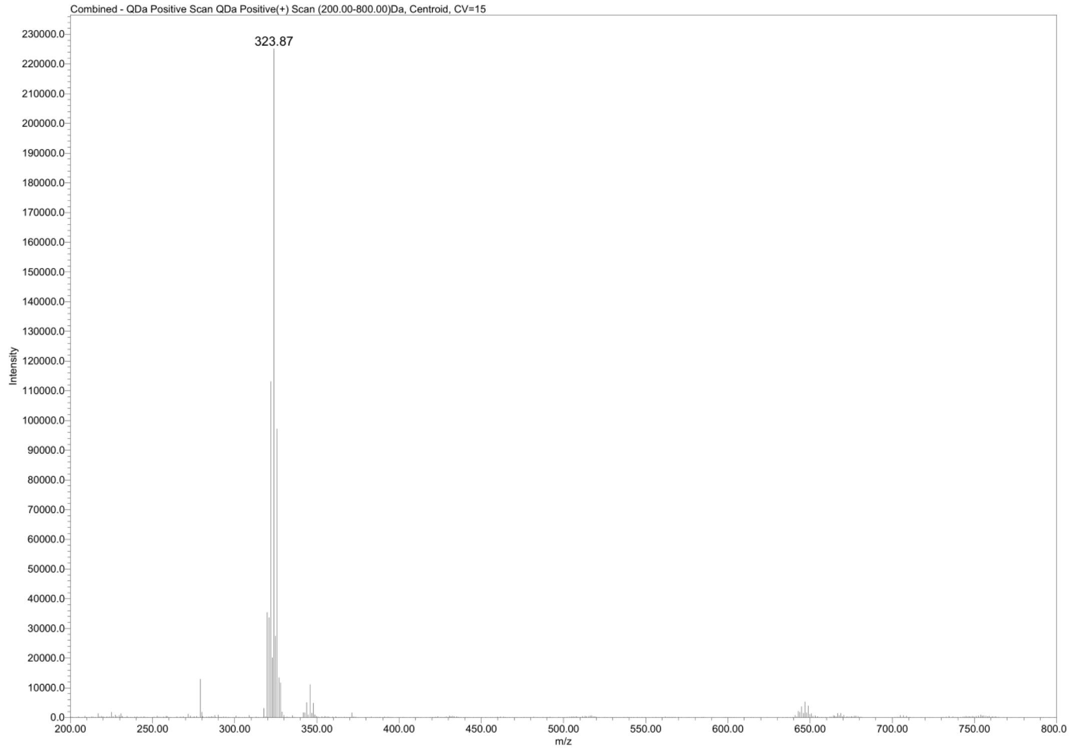

**Compound 20**

**2-(5-chloro-2-methylphenyl)-benzisoselenazol-3(2H)-one**

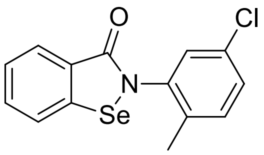


(m/z $_{calcd}$ =323.9689; m/z $_{found}$ =323.88)

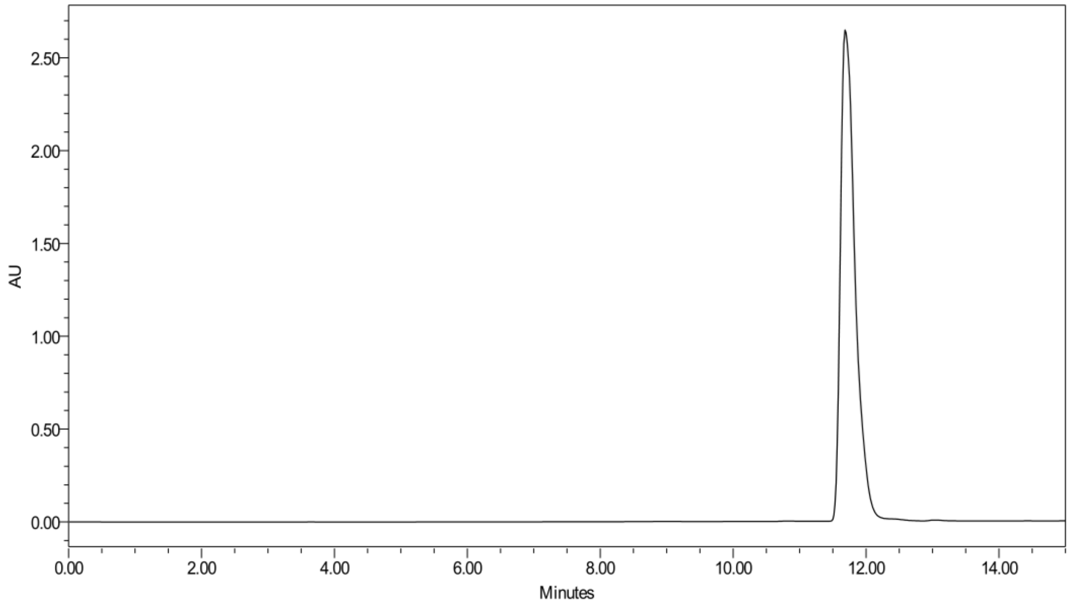


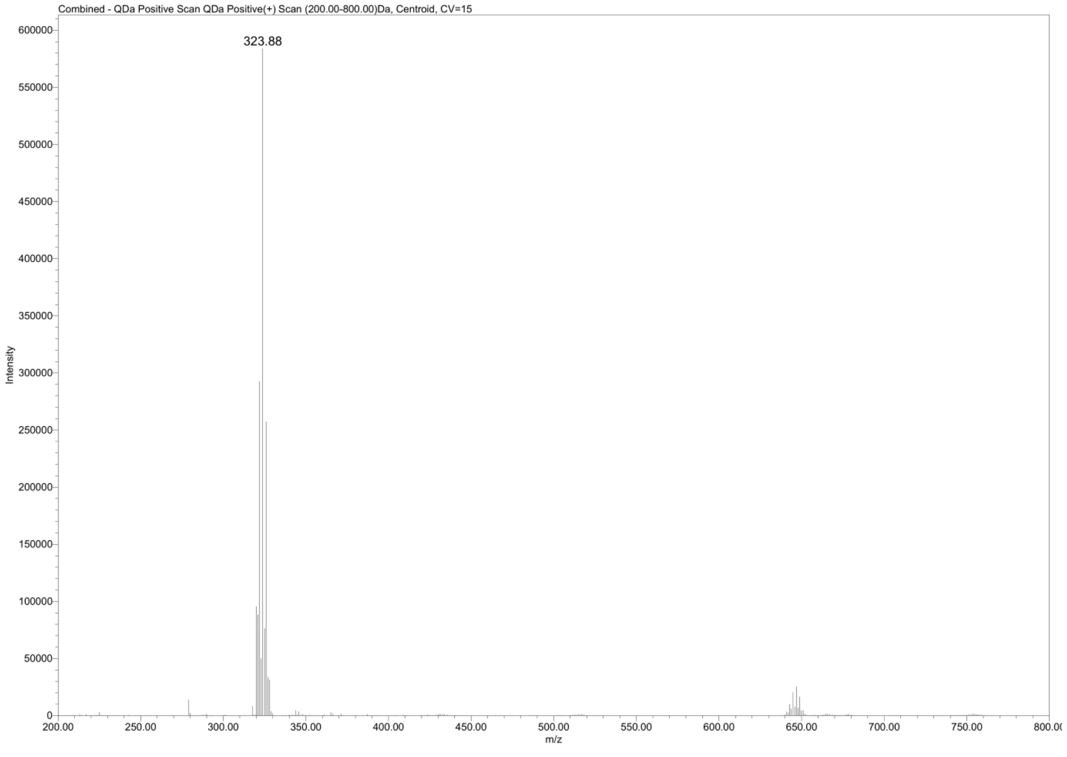

**Compound 21**

**2-(3-chloro-2-methylphenyl)-benzisoselenazol-3(2H)-one**

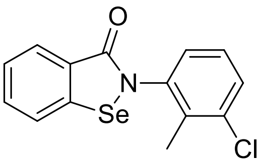


(m/z $_{calcd}$ = 323.9689; m/z $_{found}$ = 323.83)

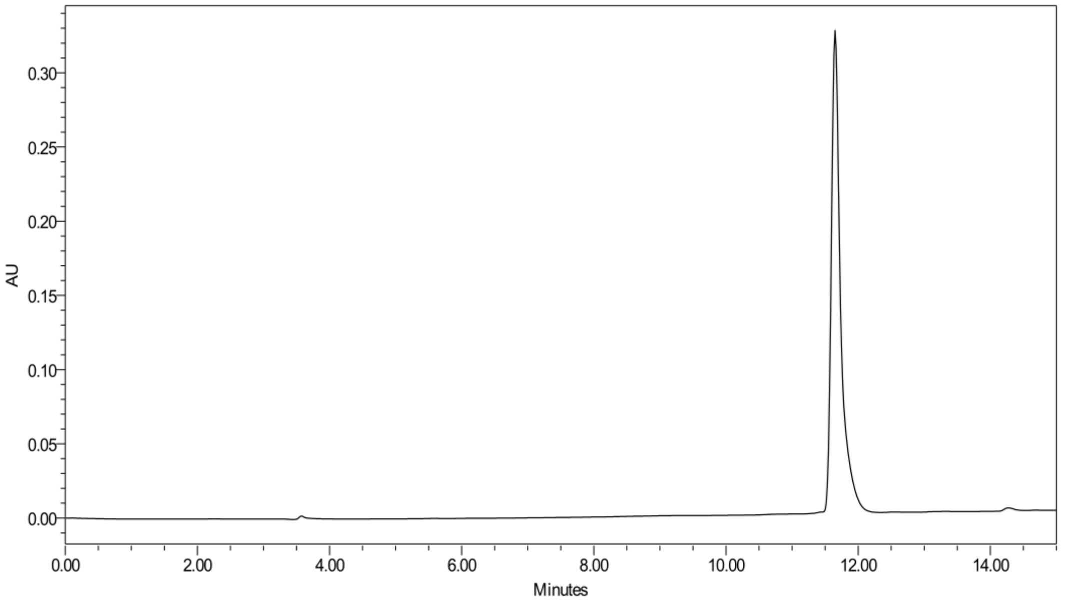


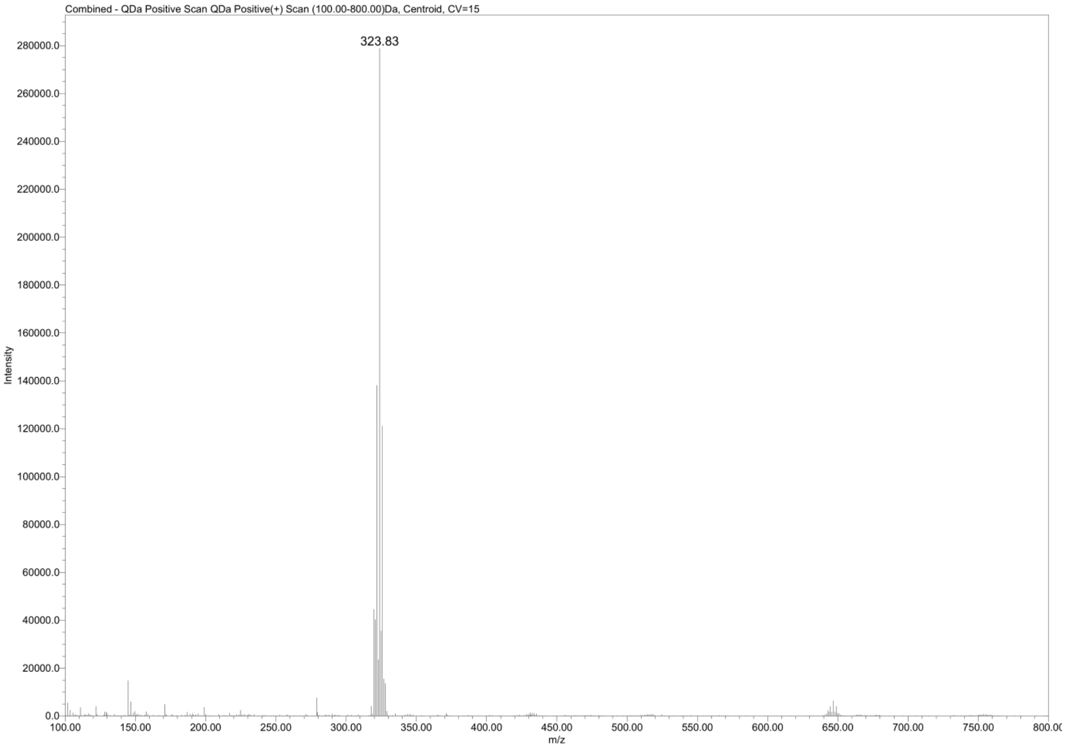

**Compound 22**

**2-(4-chloro-3-methylphenyl)-benzisoselenazol-3(2H)-one**

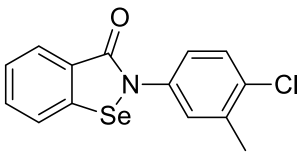


(m/z $_{calcd}$ = 323.9689; m/z $_{found}$ = 323.83)

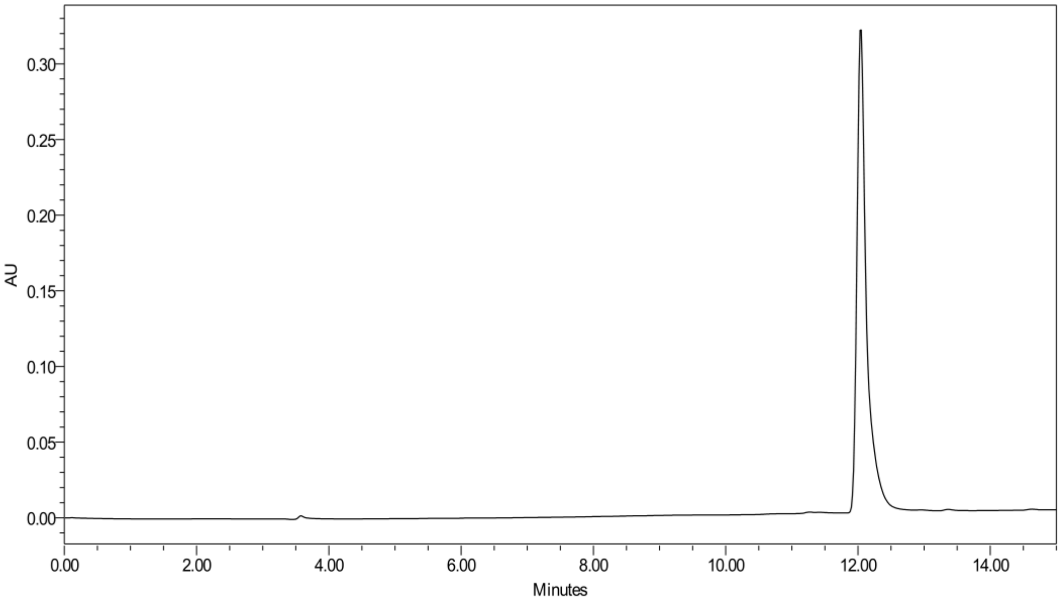


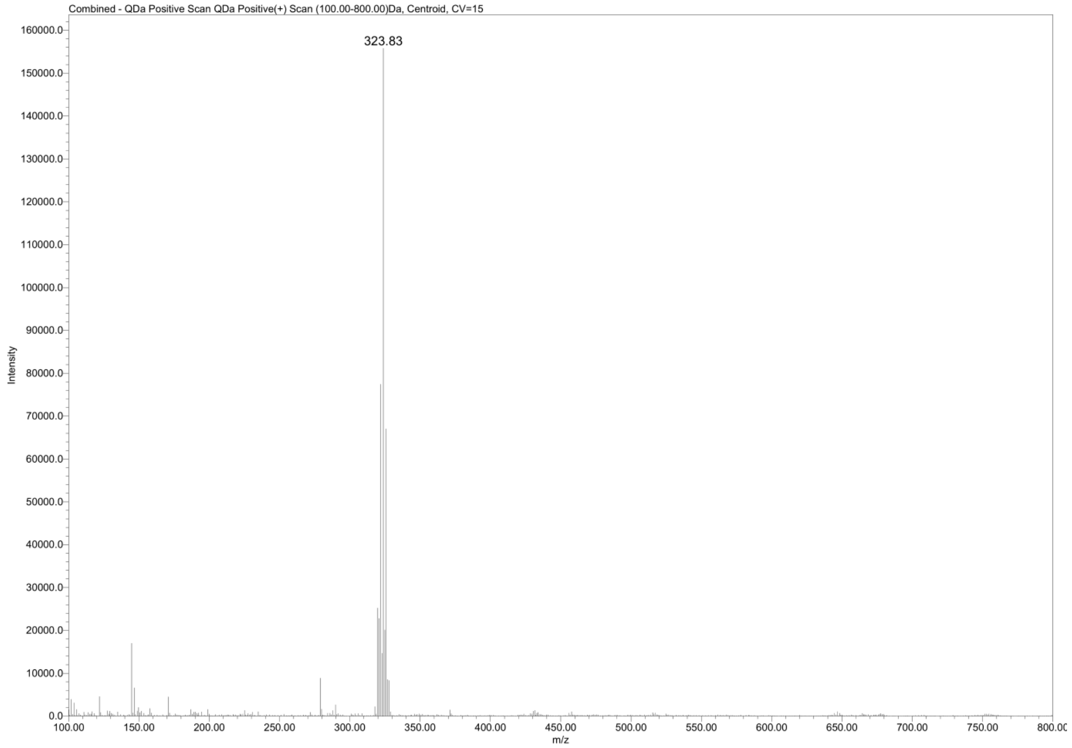

**Compound 23**

**2-(3,4-dimethoxyphenyl)-benzisoselenazol-3(2H)-one**

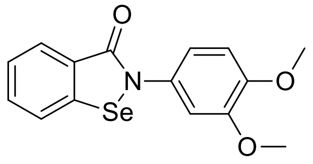


(m/z $_{calcd}$ = 336.0134; m/z $_{found}$ = 335.87)

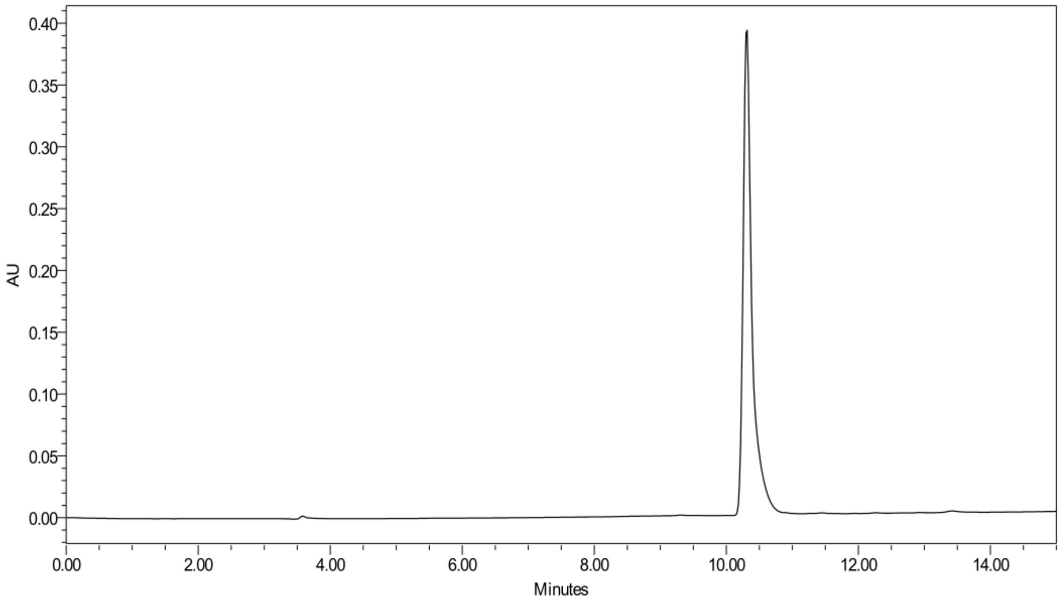


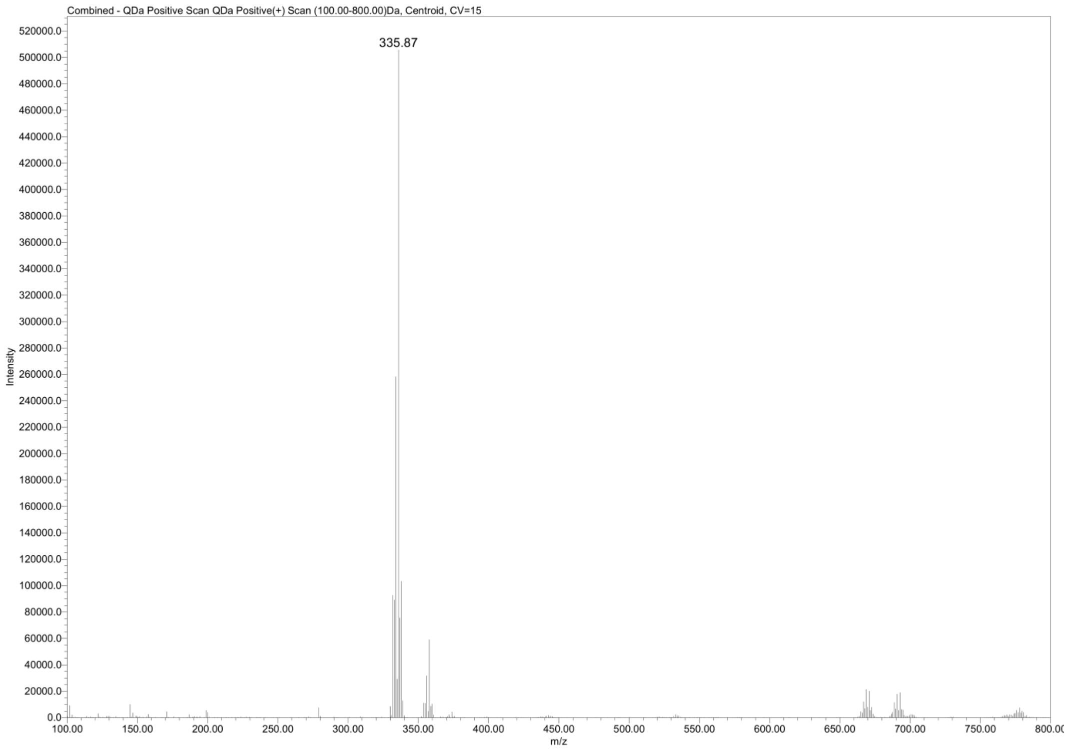
